# Peptide Design through Binding Interface Mimicry

**DOI:** 10.1101/2025.01.21.634006

**Authors:** Xiangzhe Kong, Rui Jiao, Haowei Lin, Ruihan Guo, Wenbing Huang, Wei-Ying Ma, Zihua Wang, Yang Liu, Jianzhu Ma

## Abstract

Peptides offer distinct advantages for targeted therapy, including oral bioavailability, cellular permeability, and high specificity, which set them apart from conventional small molecules and biologics. In this work, we developed an AI algorithm, named PepMimic, to transform a known protein receptor or an existing antibody of a target into a short peptide drug by mimicking the binding interfaces between targets and known binders. The structural root mean square deviation and interface DockQ with reference binders were 61% and 75% better than the best existing methods on the PepBench datasets. We then applied this novel peptide-design methodology to five drug targets: PD-L1, CD38, BCMA, HER2, and CD4. SPRi results show that 8% of the peptides exhibited dissociation constant (K_D_) values at the 10^-8^M level, and 26 peptides achieving K_D_ values as low as 10^-9^M. This success rate was 20,000 times higher than that observed in a random library screening conducted under identical conditions. PepMimic was applied to target proteins lacking available binders by first utilizing AI algorithms to design protein binders, followed by the generation of peptides through simulation of these artificial interfaces. The top-ranked peptides underwent extensive cellular validation and *in vivo* testing through tail vein injections in breast, myeloma, and lung tumor mouse models. Experimental results demonstrated effective membrane binding and highlighted the strong potential of these peptides for clinical diagnostic imaging and targeted therapeutic applications.

## Introduction

Since the advent of insulin nearly a century ago, over 80 peptide-based therapeutics have been approved for the treatment of various diseases, including diabetes, cancer, osteoporosis, multiple sclerosis, HIV infection, and chronic pain^35, 46^. The primary advantage of peptide-based therapeutics is their capacity to deliver high specificity and potency, enabling precise modulation of biological pathways. Compared to traditional small molecules and antibodies, peptide drugs tend to exhibit superior safety and therapeutic profiles due to lower toxicity, enhanced cellular permeability, and the potential for oral administration ^11, 28, 45^. One of the key observations for peptide design is that protein targets often have known binders, either naturally occurring or identified through high-throughput screening experiments from large libraries or animal immunizations. Moreover, machine learning algorithms can also be adopted to design mini-binders with high binding affinity to the protein targets^12, 47^. These existing binders provide valuable interaction templates for the design of functional peptides. On the other hand, medicinal chemists have a long history of developing peptide drugs by emulating the natural binders of target proteins. For instance, Bivalirudin^41^ is a 20-amino-acid peptide that mimics hirudin, an intrinsic occurring direct thrombin inhibitor. It binds thrombin at two sites, which are interconnected by a cleavable polyglycine chain that allows simultaneous binding of fibrinogen. Ziconotide (Prialt)^34^ is a peptide that mimics the naturally occurring cone snail venom peptide, targeting neuronal calcium channels to relieve severe chronic pain. Similarly, Leuprolide^48^ mimics GnRH, which is capable of binding to the GnRH receptor (GnRHR). Helical peptides mimicking hACE2 have also been investigated as potential inhibitors of SARS-CoV-2 by blocking its spike glycoprotein^23^. Recent literature^4^ have further identified lasso peptide, a special modality of peptide, as a potentially new direction for therapeutics by mimicking the function of natural MccJ25 on inhibiting ↵v;38. Peptide mimics of specific protein-protein interactions could also facilitate studies on modulating cellular functions by activating biological mechanisms^6, 16^. Thus, designing peptides that mimic existing binders emerges naturally as a compelling strategy for identifying candidates with desired functionality^37^. Nevertheless, there are two main drawbacks in the current methodology of designing such peptide mimics. First, designing such peptide mimics demands professional expertise and a profound understanding of interaction patterns among amino acids^15, 23^. This process often involves manually extracting binding motifs or identifying anchor amino acids at the interface through experimental methods. Second, medicinal chemists are accustomed to mimicking continuous amino acid sequences on the interface, and struggle to address discontinuous binding surfaces^5, 37^, which significantly reduces the utilization of the interface of known binders.

In this paper, we present a novel AI-based methodology, PepMimic, which can achieve sequence and structure co-design of all-atom peptide binders conditioned on a protein target and another known binder. This model facilitates the automatic creation of interface-mimicking peptides, even when the interfaces involve discontinuous amino acids on the sequence (Fig. 1a). As a geometric deep generative model, PepMimic leverages a latent space encoded by an all-atom encoder to accomplish efficient generation of peptide binders via latent diffusion process (Fig. 1b). A large-scale augmentation dataset (Methods) is introduced for pre-training before fine-tuning on high-quality protein-peptide complexes to encourage higher robustness. Leveraging the guidance of an interface encoder (Fig. 1c), which learns the interaction patterns in the well-formed latent space (Fig. 1d), PepMimic is capable of designing interface-mimicking peptides faithful to the reference binding interfaces. PepMimic effectively addressed the current challenges of designing peptide mimics by automatically learning the amino acids patterns involved in the interface interactions and representing them in a latent space. The guidance in the latent space instructs the model to connect interacting amino acids into a single peptide which can form a desired structure with a feasible sequence. Such procedure endows the model with generalizability across different interfaces to be mimicked, avoiding the need for retraining with new references. Furthermore, our model implemented a novel all-atom paradigm for peptide design. Existing models either first generate backbones and then conduct inverse folding ^17, 30, 47^ or co-designing the backbone and sequence together, and subsequently relying on third-party softwares to add side chain atoms^19, 24, 33^. These models can be recognized as modeling the conditional probability between amino acid types and 3D backbone structures, thus overlooks the importance of side-chain geometry, which plays a crucial role in defining amino acid interactions. Side-chain atoms and their spatial positioning are key to forming interactions such as hydrogen bonds and *π*-*π* stacking, making all-atom modeling essential for accurately capturing these geometric constraints and determining corresponding amino acid types. Although recent breakthroughs like AlphaFold^1, 20^ and RosettaFold^3, 26^ have enabled high-resolution structure prediction, adapting them for generative modeling, especially in the context of interface-mimicking peptide design, remains a significant challenge. Our method directly addressed this gap by capturing the joint distribution of amino acid types and all-atom structures within a latent space, offering a more precise and effective approach to all-atom peptide design, particularly for interface-mimicking peptide design.

**Figure 1.**
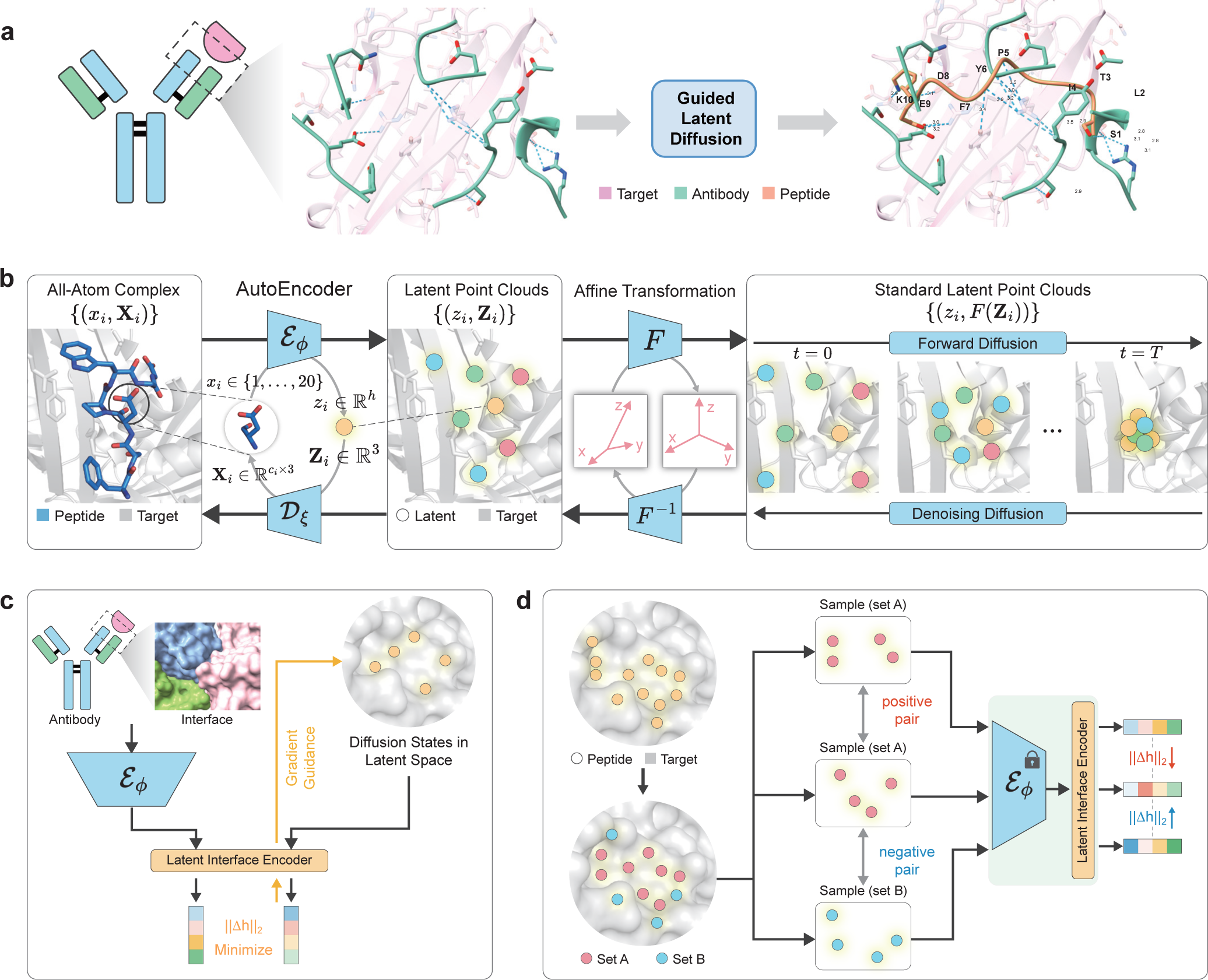
Schematic diagram of the PepMimic algorithm. Given a target protein and a specific binder, PepMimic designs interface-mimicking peptides by integrating three core modules: an all-atom autoencoder, a latent diffusion model, and a latent interface encoder. **a**, The interface of the binder to mimic (e.g. an antibody) is extracted to guide the diffusion process in the latent space. The designed peptide binder emulates the interface by automatically identifying and linearly connecting key interactions. **b**, The generative components of PepMimic incorporate an all-atom autoencoder and a latent diffusion model. Intially, the autoencoder establishes a reversible mapping from all-atom geometries to residue-level low-dimensional latent representations. Subsequently, a diffusion model is trained within the latent space to simultaneously generate the E(3)-invariant features and E(3)-equivariant features, forming a latent point cloud. Ultimately, residue types and all-atom geometries of the peptide are decoded from the invariant and the equivariant features via the autoencoder, respectively. **c**, The extracted interface is transformed into a vector-form representation by the latent interface encoder, which enables gradient guidance to minimize the distance between the given interface and the generated counterpart in the latent space. **d**, The dataset for training the latent interface encoder is constructed by downsampling protein-peptide interfaces. Peptide residues are randomly split into two non-overlapping sets, A and B. Set A is downsampled twice to form a positive pair due to their overlapping amino acids, while set B is downsampled once to derive a negative sample. The training objective is to minimize the distances between the representations of the postive pair and maximize that of the negative pair.

We first conducted *in silico* evaluations without interface guidance on 93 testing protein-peptide complexes to compare our model with state-of-the-art methods including dyMEAN^25^, HSRN^18^, and RFDiffusion^47^. Computational metrics, involving Rosetta interface energy^2^, Amino Acid Recovery (AAR), and structural Root Mean Square Deviation (RMSD) (Methods), demonstrated the superiority of our model in terms of better recovering the reference binder and generating candidates with lower energy. Next, we designed 384 peptides for five drug targets, PD-L1, CD38, BCMA, HER2, and CD4, by mimicking their natural receptors, antibodies, and nanobodies. 8% peptides achieved a binding affinity at or above the level of 10^-8^M, with 26 peptides reaching a binding affinity of 10^-9^M. For targets without known binders, our model can simulate binding interfaces of mini-binders generated by existing AI systems. In this way, we introduce a novel design paradigm, which involves initially designing protein binders and then using these mini-binders as templates for the design of short peptides. We designed 384 peptides to mimic interfaces formed by the helical protein binders designed by RFDiffusion for TROP2 and CD38, among which 14 peptides reached a binding affinity at or stronger than the level of 10^-8^M. For PD-L1, CD38, HER2, and TROP2, we further selected designed peptides with high affinities to perform more rigorous validation of their specificity and affinity at the cellular level, each showing effective binding to the cell membrane and distinct binding contrasts between positive and negative cells. Their effectiveness as molecular targeting probes was further validated by tail vein injections in breast and pancreatic tumor mouse models. Significant accumulation at tumor sites was observed within 0.5 hours after post-injection, highlighting their promise for clinical diagnostic imaging and targeted therapy. These experimental results validated the ability of PepMimic to design peptide mimicries for given interfaces, posing significant potential to facilitate the discovery of functional peptides in biological researches and therapeutic applications.

## Results

### Overview of PepMimic

PepMimic is composed of three core modules: an all-atom autoencoder, a latent diffusion model, and an interface encoder for guided mimicry design (Fig. 1a). PepMimic was trained to co-design the sequence and the all-atom structure of peptide binders at specific binding sites (Fig. 1b). The all-atom encoder compresses all the atoms associated with each amino acid into a single embedding, which establishes a reversible mapping from the all-atom geometries of peptides to the latent point clouds at the amino acid level. The training loss of the encoder is to reconstruct the types and atomic coordinates of each amino acid based on its latent embedding. Next, a probabilistic diffusion model was trained to gradually transform noises from standard Gaussian into a suitable latent point cloud conditioned on the binding site. During generation, the latent point cloud is first sampled from the latent space with Gaussian noise, transformed by the diffusion model, and then decoded into an all-atom peptide structure via the decoder module.

To mimic the binders, the same encoder function was adopted to project the all-atom geometry of the reference binding interface into the same low-dimensional space as a latent point cloud. Based on these point clouds, an interface encoder further represent each point cloud as one single numerical vector. The reference binder and the peptide being represented each correspond to one vector and their distance in vector space reveals the similarity of their binding patterns with the same protein target (Fig. 1c). Therefore, we could rely on the distance between these two vectors to guide the generative diffusion process via gradient descend, gradually aligning the latent point cloud with the desired interface in the latent space. To achieve effective guidance, training of the latent interface encoder aims to endow the representation with the capability of distinguishing similar and dissimilar interfaces (Fig. 1d) by contrastive learning.

### Performance on target-specific peptide design

To assess the peptide binder design capability of PepMimic, we conducted evaluations on a non-redundant test set comprising 93 curated protein complexes from the literature^43^. Complexes with target proteins exhibiting over 40% sequence identity determined by mmseqs2^39^ were excluded from the training and the validation sets to ensure the generalizability across different targets. For each target protein, we sampled 40 candidates for all the baseline models. The generated 3D peptides were evaluated based on the following metrics (Methods): 1) Rosetta interface energy^2^(Δ*G*), Root Mean Square Deviations (RMSD) of C_*α*_ coordinates, DockQ for the all-atom interface, and Amino Acid Recovery (AAR). Except for Δ*G*, all metrics were computed by comparison to known peptide binders in the test set. We compared PepMimic to the state-of-the-art protein design models including RFDiffusion^47^ and two sequence-structure co-design models for antibody CDR loops, HSRN^18^ and dyMEAN^25^, which were both retrained on protein-peptide complexes, the same as our model. For RFDiffusion, we directly used its official weights and pipeline for binder design since it did not open-sourced the codes for training. Technical details relating to the implementation of baseline methods could be found in the Supplementary Notes 2. For fair comparison, candidates generated by all models endured fast relax via Rosetta only once before calculating the Δ*G*.

PepMimic consistently outperforms all baselines across all metrics, regardless of peptide length (Fig. 2a). The recovery rate of the reference peptides tends to decrease as the peptide length increases, due to the larger combinatorial search space of amino acids. Rosetta Δ*G*, opposite to the trend, is lower for longer peptides, as they usually form larger interface with lower interface energy (Supplementary Fig. 1). Evaluations categorized by different secondary structures further confirmed the superiority of PepMimic over existing baselines (Fig. 2b). In contrast to RFDiffusion, PepMimic achieved higher AAR (40% vs 33%, *p* ≤ 0.05) on peptides without secondary structures, which is expected, as RFDiffusion is pre-trained on general proteins and may be better suited for those with stable secondary structures^47^. The trending of metrics with enlarged sample sizes indicated gradual saturation beyond 30 samples (Fig. 2c). Therefore, a sampling pool with above 30 generations are sufficient to produce at least one satisfactory candidate which approximates the performance upper bound of the model. HSRN, while achieving high AAR through autoregressive sequence modeling, struggles with generating peptides with accurate binding structures. DyMEAN, on the other hand, shows early metric saturation with just a few samples, due to a lack of diversity (Fig. 2c).

**Figure 2.**
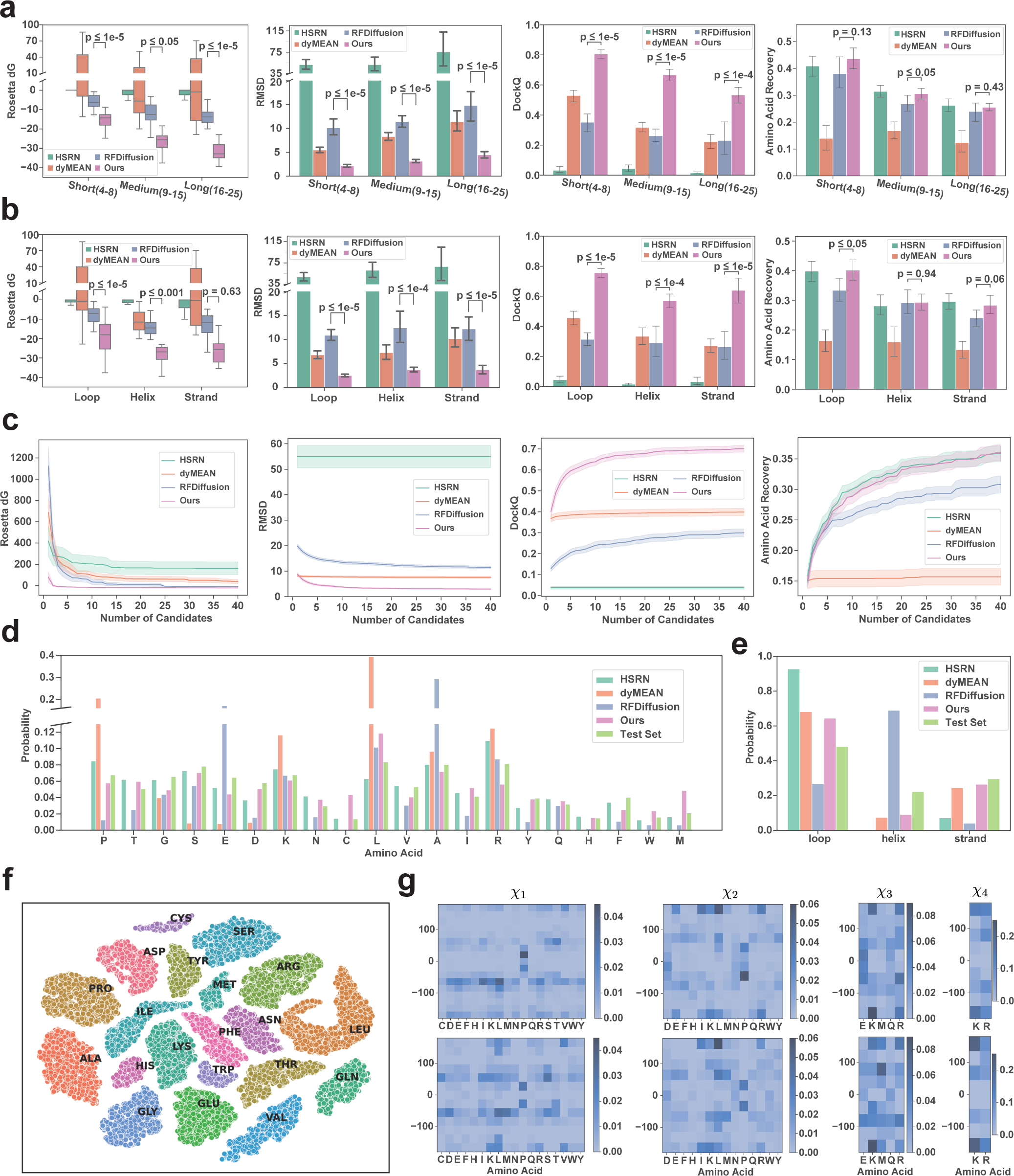
Evaluations on target-specific peptide design on a non-redundant test set collected from literature. Basic metrics include Rosetta interface energy (Δ*G*), Root Mean Square Deviation (RMSD)of C_*α*_ coordinates, full-atom DockQ, and Amino Acid Recovery (AAR). **a**, Metrics of different models on the test set categorized by the lengths of the peptides. **b**, Metrics of different models on the test set categorized by the secondary structure of the reference peptides. **c**, Saturation trend of metrics when expanding the number of generated candidates. **d**, Probability distribution of generated amino acid types for different models compared to the reference distribution. e, Probability distribution of secondary structures in the generated peptides compared to the reference distribution. **f**, T-SNE visualization of amino acid representations in the latent space, with an Adjusted Rand Index (ARI) of 1.0 for amino acid categories. The representation distances have a Pearson correlation of 0.47 to BLOSUM62 matrix (p-value< 0.001). **g**, Distribution of side-chain torsional angles from *χ*_1_ to *χ*_4_. The top row shows the reference distribution from the test set, while the bottom row shows the distribution from the peptides generated by PepMimic.

Next, we compared the amino acid and secondary structure distributions of the peptides generated by different models and those in the test set, in order to investigate whether a particular model exhibits a preference for producing specific amino acids or secondary structures. The amino acid distribution generated by PepMimic is more similar to the test set (KL=0.06), while dyMEAN (KL=0.89) and RFDiffusion (KL=0.38) exhibited a higher propensity for generating proline, glumatic acid, lysine, and alanine (Fig. 2d). The distribution of secondary structures generated by PepMimic leaned slightly towards loops, whereas RFDiffusion showed a strong preference for generating helical peptides, which is consistent to the literature^47^(Fig. 2e). T-SNE^44^ visualization of the latent amino acids representations demonstrated a continuous and well-formed latent space for the diffusion of sequences (Fig. 2f). The representations were closely clustered to distinguish different amino acids (ARI^40^= 1.0). In addition, amino acids with similar physicochemical properties and biological functions were proximally located in the latent space made by the all-atom autoencoder. For instance, three amino acids with polar uncharged side chains, asparagine, threonine, and glutamine, were closely located in the latent space. Three amino acids with aromatic side chains, tyrosine, phenylalanine, and tryptophan, were also adjacent in the latent space. Distances of amino acids in the latent space moderately aligned with the BLOSUM62 substitution scores^14^ (Pearson correlation= 0.47, *p*-value< 0.001). Such a similarity-informed latent space facilitates diffusion to generate accurate or functionally similar amino acids. The generated distribution of side-chain torsional angles (x_1_ to *χ*_4_, bottom row in Fig. 2g) matched well with that of the test set (top row in Fig. 2g), indicating a capability of generating accurate all-atom geometries by PepMimic. Overall, extensive evaluations above demonstrated that PepMimic consistently outperformed state-of-the-art models regarding co-designing the sequence and the all-atom structures of peptide binders given specific binding sites, as evidenced by comprehensive *in silico* metrics.

### Performance of interface-mimicking

Next, we assessed the efficacy of interface-mimicking generation under the proposed latent guidance. We first derived a computational metric, named **Interface Hit**, to measure quality of the alignment between the designed interface and the interface to be mimicked (Fig. 3a). For a pair of 3D interfaces, where the target protein has a binding site comprising *N* amino acids, the distance from one amino acid on the binder to another amino acid on the binding site are calculated, which yields an *N*-dimensional feature vector for each amino acid. Then for each pair of amino acids on the two binders, we calculated their Pearson correlation *R* of feature vectors. Amino acid pairs with *R* > 0.9 and BLOSUM62 score above zero were identified as a pair of ‘Interface Hit’, indicating they share a certain degree of geometric and functional similarity.

**Figure 3.**
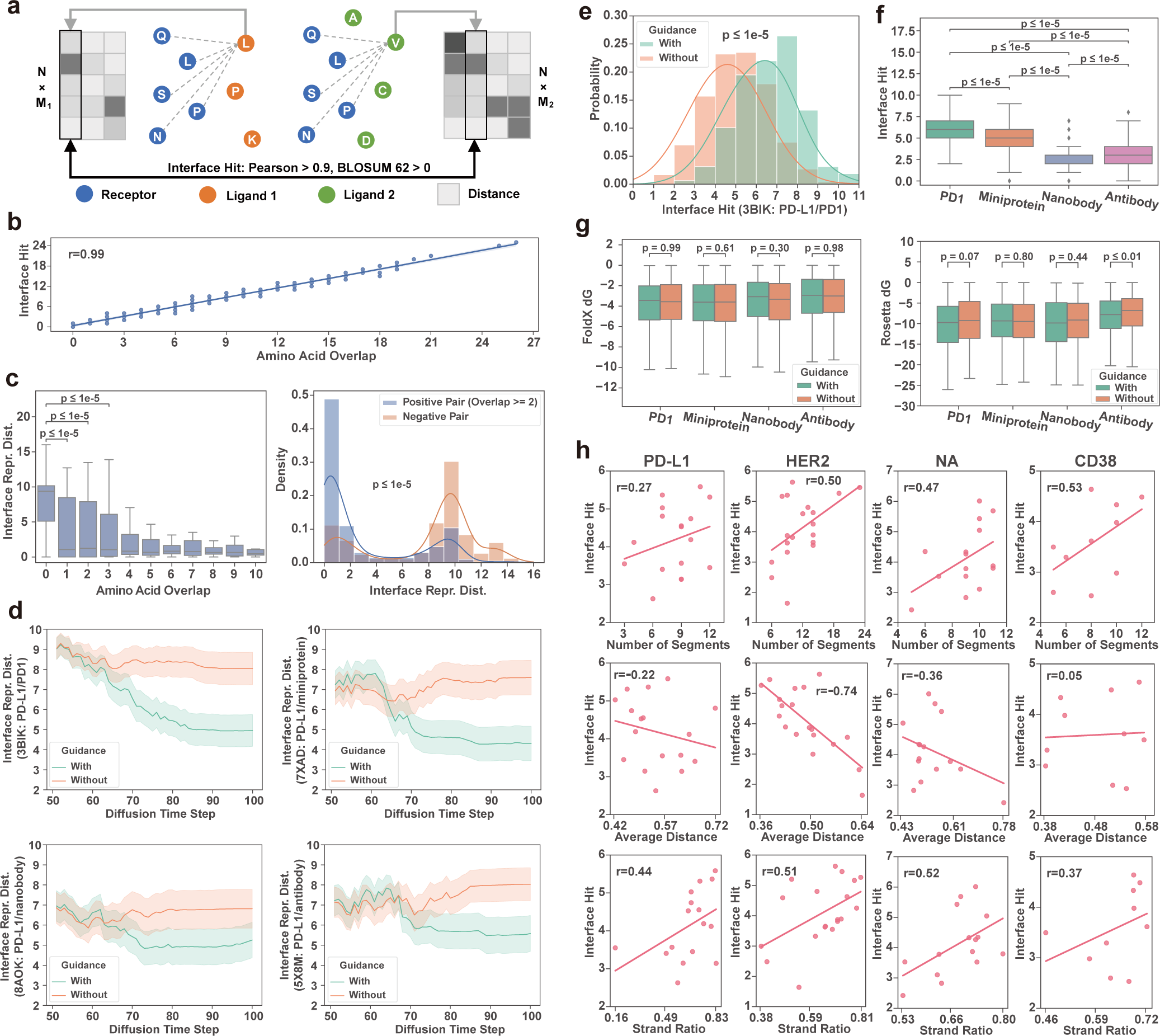
Evaluations on the latent interface encoder and analyses on Interface-Mimicking Capability of PepMimic. **a**, Definition of interface hit, which quantitatively measure the similarity of two interfaces. The binding site on the target protein comprises *N* residues, whereas two ligands comprises *M*_1_ and *M*_2_ residues, respectively. The interface hit metric counts the number of hit pairs on the two interfaces, where “hit” is defined as residue pairs on the two ligands with a Pearson correlation of the distances to *N* target residues above 0.9 and a BLOSUM62 substitution score above 0. **b**, Pearson correlation between interface hits and known amino acid overlaps on the dataset constructed via the downsampling strategy (p < 0.001). **c**, Distances of representations given by the latent interface encoder categorized by number of amino acid overlaps in the interface pairs (left). Comparison of representation distance distribution of postive pairs (with overlaps above 2) and negative pairs (without overlap). **d**, The trajectories of representation distances between the reference interface and the interface formed by the generated peptide under latent guidance. Results on four types of reference ligands on PD-L1 are visualized: the natural receptor PD1, AI-designed miniprotein, nanobdy, and the antibody durvalumab. **e**, Comparison of distribution of interface hits with and without guidance. **f**, Comparison of interface hits on different types of ligands for PD-L1. **g**, Comparison of FoldX Δ*G* and Rosetta Δ*G* for generated peptides with and without guidance, categorized by different type of reference interface. **h**, Pearson correlation between interface characteristics and interface hits of generated peptides on four target proteins: PD-L1, HER2, viral neuraminidase (NA), and CD38. The characteristics include number of segments, average distances between residues, and strand ratio on the reference interface.

To verify the fidelity of the proposed interface hit, we down-sampled the interfaces in the test set to construct interface pairs with known alignments of amino acids (Fig. 1d, Supplementary Notes 1.2.1). Interface hit of these test pairs exhibited high correlation with the number of overlapping amino acids (Pearson correlation= 0.99, P < 0.001), indicating its reliability in measuring the alignment of the interfaces. To evaluate our ability to accurately mimic a given interface, we initially assessed the PepMimic’s capacity to differentiate between similar and dissimilar interfaces. This was achieved using pairs in the test set that were constructed through interface partitioning and the down-sampling strategy (Supplementary Notes 1.2.2). The distances between latent representations of the interface pairs decreased as the number of overlapping amino acids increased, with even a single pair of overlapping amino acids sufficient to significantly (*p* ≤ 10^-5^) reduce the distances (Fig. 3c, left). The encoder effectively distinguished positive pairs (with more than two overlapping amino acid pairs) from negative pairs (with no overlaps) with significant (*p* ≤ 10^-5^) distribution difference in representation distance, establishing it as a reliable differentiable proxy for approximating interface similarity (Fig. 3c, right).

Next, we evaluated the effectiveness of gradient-based guidance in aligning the latent representations of generated peptides with the reference interfaces (Fig. 3d). Experiments were conducted on PD-L1 interfaces with four binding partners: its natural receptor PD1^31^, a mini-protein binder designed by MaSIF suite^13^, a nanobody^22^, and the antibody durvalumab^29^. K_D_ values of PD1, the AI-designed mini-protein, the nanobody, and the antibody to PD-L1 are 8.2 *µ*M^7^, 374 nM^13^, 44 nM^22^, and 0.667 nM^42^, respectively. The guidance procedure significantly reduced interface representation distances compared to unguided counterparts (*P* < 0.01), particularly after 50 diffusion steps (out of 100). The distribution of interface hit for the generated peptides further confirmed the efficacy of the latent guidance (Fig. 3e, Supplementary Fig. 2a). The magnitude of distance reduction varied, with PD1 and the mini-protein showing larger gaps than the nanobody and antibody, reflecting different levels of mimicry difficulty. PD1 is a natural receptor for PD-L1 and their binding interface is optimized for efficient interaction during evolution^50^. Mini-proteins, especially those designed by machine learning algorithms, often have streamlined and specific interaction interfaces tailored for enhanced binding^8^ (Supplementary Fig. 2b). Nanobodies and antibodies generally have larger and more complex binding interfaces. They often make extensive contacts with the antigen, involving a variety of loops and secondary structures^9, 36^ (Supplementary Fig. 2b). This complexity might make it more challenging for the algorithm to accurately mimic these interfaces due to higher flexibility and intricate contact points. These findings were also supported by the distinct distributions of interface hit for the four interfaces, with higher hits for PD1 and the mini-protein indicating easier mimicry compared to the nanobody and antibody (Fig. 3f). These evaluations demonstrated that latent guidance, facilitated by the latent interface encoder, had effectively directed the latent diffusion process towards generating 3D peptides that mimicked specified interfaces. Additionally, comparisons of Δ*G* values with and without guidance showed that guided generation produced superior or comparable Δ*G*, as assessed by FoldX^38^ or Rosetta^2^, confirming that the guidance procedure maintained realistic candidate generation (Fig. 3g).

In-depth analyses were conducted to investigate the correlation between the characteristics of reference interfaces and the distribution of interface hits, aiming to identify which types of interfaces are easier to mimic *in silico*. A total of nine characteristics of the binding partners forming the reference interfaces with the target protein were analyzed, including helix length (HL), helix ratio (HR), loop length (LL), loop ratio (LR), strand length (SL), strand ratio (SR), maximum segment length (MSL), number of segments (NoS), and average distance (AD) between interface amino acids, with secondary structures classified according to the DSSP algorithm^21^ (Fig. 3h, Supplementary Fig. 3a-d). Results from four distinct target proteins: PD-L1, HER2, viral neuraminidase (NA), and CD38—revealed consistent trends: interface hits of generated peptides are negatively correlated with average distances between interface residues, and positively correlated with the number of segments and strand ratio at the interface (Fig. 3h). Smaller average distances between interface residues indicate a more compact and tighter binding interface, which facilitates the peptide mimicry design by making it easier to connect adjacent binding amino acids into one peptide. Statistics showed that the number of segments is negatively correlated with average distance between interface amino acids (Supplementary. 3e), thereby reducing the difficulty for designing a mimicking peptide. However, if the segments are scattered far apart, forming large and complex binding interfaces, designing a peptide mimicry becomes more challenging as can be seen in the antibody case for PD-L1. Strands provide linearly extended surfaces for interaction, which is a simple pattern for generative models to mimic; hence, an increased strand ratio typically leads to higher interface hits. However, special caution shold be paid to the designed peptides mimicking strands, as linear peptides rarely form such secondary structures naturally. Thus, such peptide mimicries of strands with high interface hit are likely to be artifacts, which we will demonstrate in the subsequent results of SPRi assays. Other tested characteristics showed no significant correlation or exhibited inconsistent trends across different targets (Supplementary Fig. 3a-d).

### Results on Mimicking Existing Binders for Target Proteins

We evaluated the binding affinity of the generated peptides using SPRi experiments for five targets: PD-L1, CD38, BCMA, HER2 and CD4 (Methods). For each target, we collected protein complexes with solved 3D structures which include both targets and binders (Supplementary Table 1-5). The target binders included antibodies, nanobodies, and natural receptors to the targets, which were later used for interface-mimicking design for our algorithm. Designed peptides were ranked by Rosetta^2^, FoldX^38^, and AlphaFold Multimer^10, 20^, and the top 384 candidates were selected for the next SPRi experiments (Methods). Successful hits were characterized by a dissociation constant (K_D_) below 100 nM. Our model achieved success rates above 10% for targets CD38, HER2, and CD4, while PD-L1 and BCMA showed relatively lower success rates (Fig. 4a). Success rates for mimicking antibodies and natural receptors were approximately 2% higher than for nanobodies, with a similar number of candidates tested for each binder type (Fig. 4b). Furthermore, for the successful hits, while the structures displayed certain clusters, the sequence exhibited much higher diversity (Supplementary Fig. 4).

**Figure 4.**
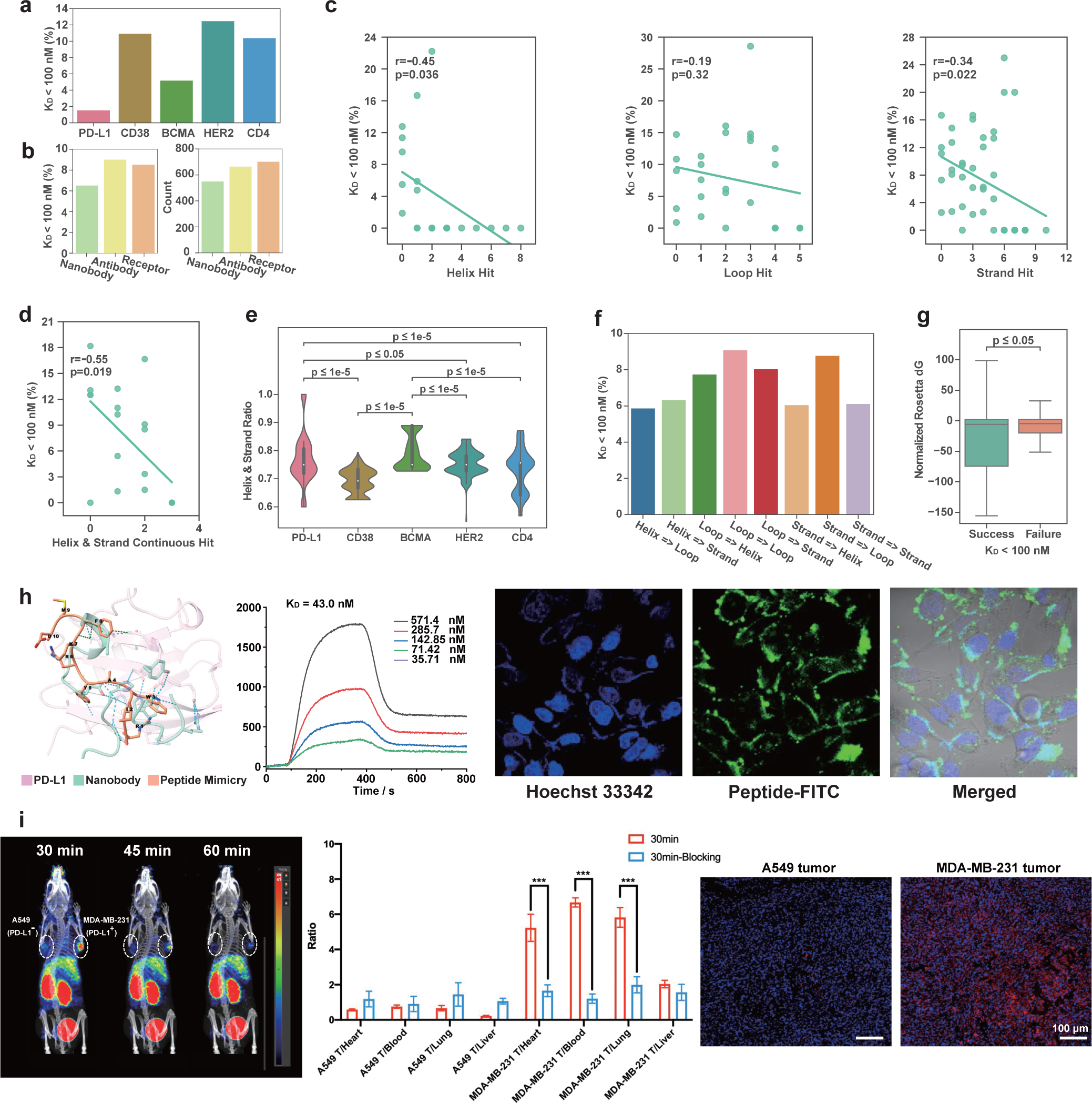
Results and analysis of molecular SPRi experiments, cellular assays, and *in vivo* validations for peptide mimicry. **a**, Success rates (K_D_ < 100 nM) across different target proteins. **b**, Success rates and the number of experimentally tested candidates categorized by the type of reference binder for mimicry. **c**, Correlation between success rate and the number of interface hits with secondary structures (helix, strand, or loop) on the reference binder. The hits were identified using the previously described “interface hit” algorithm in this paper. **d**, Correlation between success rate and the number of continuous hits on helices and strands from the reference binder. **e**, Proportion of helices and strands in the reference binder, categorized by target protein. **f**, Success rates categorized by the type of secondary structure mapping between the mimicry and the reference binder (e.g., Helix=>Loop indicates peptides using helices to mimic loops on the reference interface). **g**, Comparison of Rosetta interface energy between successful and failed candidates, with energy values normalized by the average energy of candidates mimicking the same reference binder. **h**, SPRi and confocal imaging analysis on H1975 cells (PD-L1 positive) for the specificity of the selected peptide for human PD-L1 *in vitro*. Scale bar: 20 *µ*m. **i**, Micro PET images obtained at 30, 45, and 60 min post injection of ^68^Ga– DOTA – RIWAYRKFMD in mice with bilateral A549 and MDA-MB-231 tumors xenografts (n=5). Tumor-to-normal tissue uptake ratio in MDA-MB-231 and A549 tumor-bearing mice. Immunofluorescence staining displaying PD-L1 expression in tumor tissues.

Based on the experimental results, we investigated which types of interaction interfaces are more easily to induce successful peptide mimicries. First, we observed there was a moderate negative correlation between success rates and the number of interface hits on helices or strands (Fig. 4c, p < 0.05), while no significant correlation was found for hits on loops (Fig. 4c, p = 0.32). We reasoned that this was because the conformation flexibility in linear peptides made it difficult to form stable secondary structures like helices and strands, which was supported by a more negative correlation between success rates and hits on continuous helices or strands (Fig. 4d, p < 0.05). While the *in silico* analyses found that interface hits were positively correlated with strand ratio (Fig. 3h), SPRi results indicated these interface hits on strands might be artifacts. It was highly probable that the true conformation of the designed peptides might not adopt the stable strands in the absence of contextual assistance from surrounding strands. Such phenomena also explained the relatively lower success rates on PD-L1 and BCMA, whose reference binders possessed more helices and strands on the binding interface compared to other targets (Fig. 4e, p < 0.05), making it more challenging to design successful peptide mimicries. Besides, PD-L1 was also known to be difficult to find small binders due to the flat binding site^13, 49^. Additional statistics demonstrated that the success rates were higher when peptides employed loops to mimic interface structures, or when the reference interfaces mostly consist of loop regions (Fig. 4f). Other interface properties for which no significant correlations were observed were included in Supplementary Fig. 5. Furthermore, Rosetta interface energy^2^ was found to be moderately effective in distinguishing positive binders from negative ones (Supplementary Notes 3, Fig. 4g).

We selected the peptide mimicry binding to PD-L1 with good affinity (RIWAYRKFMD, K_D_ = 43.0 nM) to conduct additional cellular assays (Fig. 4h) and further *in vivo* validation on tumor mouse models (Fig. 4i). The selected peptide was labeled with fluorescein isothiocyanate (FITC), and confocal imaging confirmed that the FITC-labeled peptide effectively binds to PD-L1-positive H1975 cells. These findings demonstrated that the candidate peptide possesses both high affinity and specificity (Fig. 4h, Supplementary Fig. 6a). To further assess its *in vivo* targeting capability for PD-L1, the selected peptide was conjugated with DOTA to chelate the radionuclide Gallium-68. Small-animal PET/CT imaging revealed a 8.85-fold greater uptake in the MDA-MB-231 tumor compared to A549 tumor (6.02 ± 0.4% ID/cc vs. 0.68 ± 0.1% ID/cc) after intravenous injection of the ^68^Ga– DOTA – RIWAYRKFMD probe 30 minutes (Fig. 4i). An examination of the delineated region of interest, encompassing the entire outer edge of the tumor, revealed the following tumor-to-blood ratios: A549 (0.75 ± 0.09) and MDA-MB-231 (6.67 ± 0.26). Over time, the majority of the ^68^Ga– DOTA – RIWAYRKFMD probe was cleared renally within 60 minutes. Consistent with PET imaging findings, immunofluorescence staining of tumor tissues confirmed that our selected peptide specifically recognized the PD-L1-positive MDA-MB-231 tumor, while showing negligible binding to the PDL1-negative A549 tumor (Fig. 4i). Collectively, these results demonstrated that the targeting peptide developed through this strategy exhibited remarkable specificity for PD-L1 both *in vitro* and *in vivo*, highlighting its potential for targeted cancer imaging. Further *in vitro* and *in vivo* validation for peptide mimicries binding to CD38 and HER2 were included in Supplementary Fig. 6b-c and 7. In addition, we also constructed a random peptide library with a capacity of 10^7^ to experimentally screen peptides targeting PD-L1, which resulted in only 8 positive peptides with a K_D_ of 10^-8^M (Methods). *In vivo* control experiments with the best peptide obtained from high-throughput experiments from a random library were included in Supplementary Fig. 8 for comparison. PepMimic achieved a success rate 20,000 times higher than that of the experimental approach in terms of enriching high-affinity peptides.

Exploring successful designs on different target proteins, we discovered diverse mimicking patterns. Typical examples included that 1) using one peptide to mimic the amino acids crossing the; 3-sheet (Fig. 5a); 2) mimic the loops between consecutive; 3-sheets (Fig. 5b); 3) one peptide connecting linearly distributed loop segments (Fig. 5c); 4) crossing two parallelly displaced; 3-sheets (Fig. 5d); 5) linking antibody CDRs (Fig. 5e), as well as bridging two separate interfaces (Fig. 5f).

**Figure 5.**
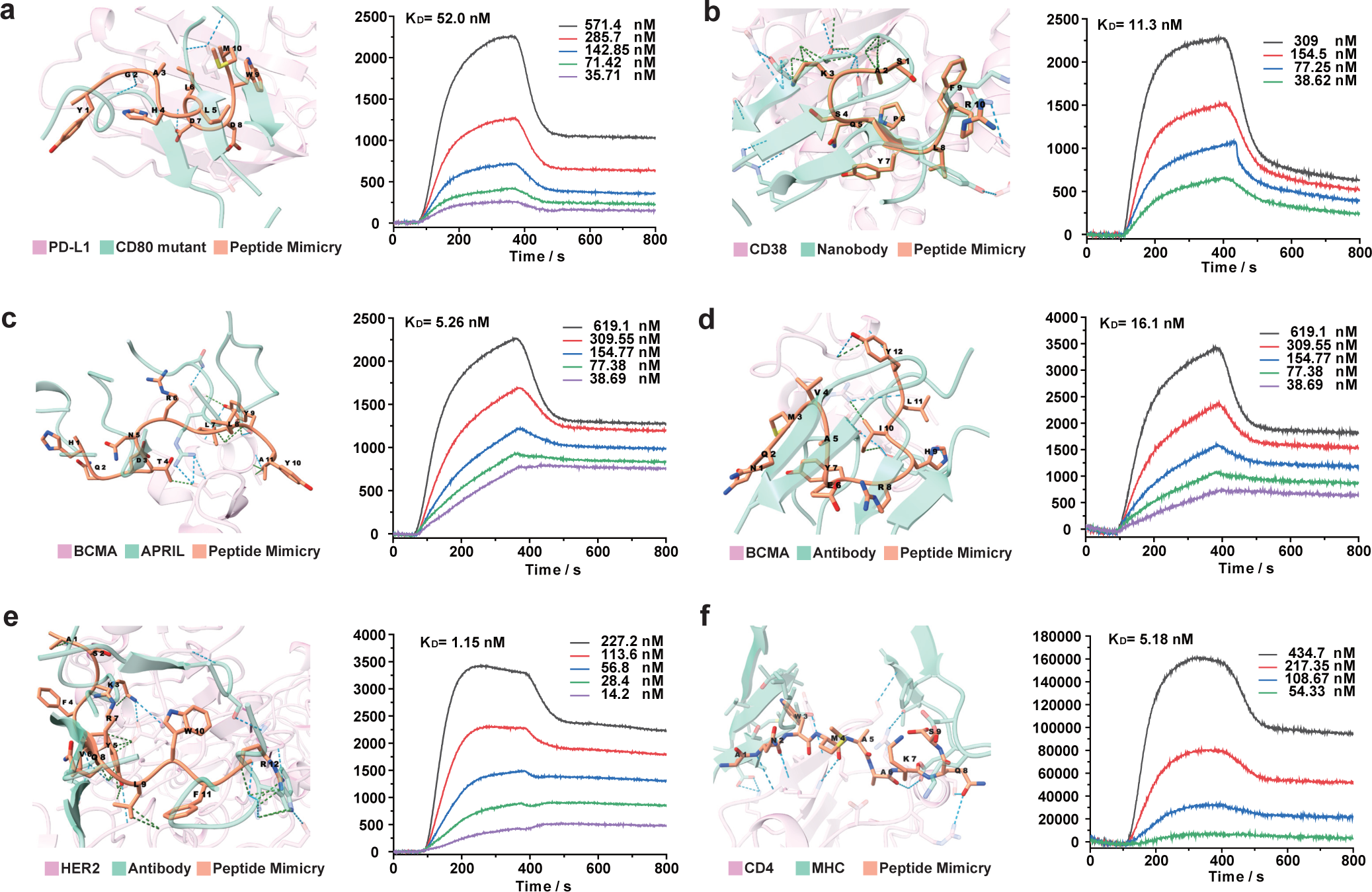
Generated structures and SPRi results of designed peptide mimicries on different target proteins. **a**, A peptide mimicry of CD80 mutant binding to PD-L1 (PDB: 7TPS) with K_D_ = 52.0 nM. **b**, A peptide mimicry of a nanobody binding to CD38 (PDB: 5F1K) with K_D_ = 11.3 nM. **c**, A peptide mimicry of APRIL binding to BCMA (PDB: 1XU2) with K_D_ = 5.26 nM. **d**, A peptide mimicry of an antibody binding to BCMA (PDB: 4ZFO) with K_D_ = 16.1 nM. **e**, A peptide mimicry of an antibody binding to HER2 (PDB: 3N85) with K_D_ = 1.15 nM. **f**, A peptide mimicry of MHC binding to CD4 (PDB: 3T0E) with K_D_ = 5.18 nM.

### Results on Mimicking AI-Generated Binders

In the previous study, we found that our method had certain advantages over RFDiffusion in terms of sequence recovery rate and structural RMSD, even without the usage of guidance (Figs. 2a-c). Next, we studied whether our algorithm could be adopted to improve RFDiffusion by mimicking the peptides and mini-binders generated by RFDiffusion (Fig. 6a). We applied this methodology to two targets: CD38 and TROP2. The epitopes were manually selected based on structural-functional analyses from existing literature^32^ (Supplementary Table 6). For each target, we generated a total of 12,000 candidates with RFDiffusion^47^, ranked them using Rosetta^2^ and FoldX^38^. The top 1,500 candidates for each of the two ranking approaches, summing up to 3,000 generations, were then subjected to co-folding with the targets using AlphaFold Multimer^10, 20^. Complexes with an average pLDDT score above 90 at the interface, including residues within 6Å C_(3_ distance to the binding partner, were chosen for peptide mimicry design (Fig. 6b). For TROP2, due to only one complex meeting the threshold, we also included the second highest scoring complex (pLDDT = 88), which was close to the threshold (Fig. 6b).

**Figure 6.**
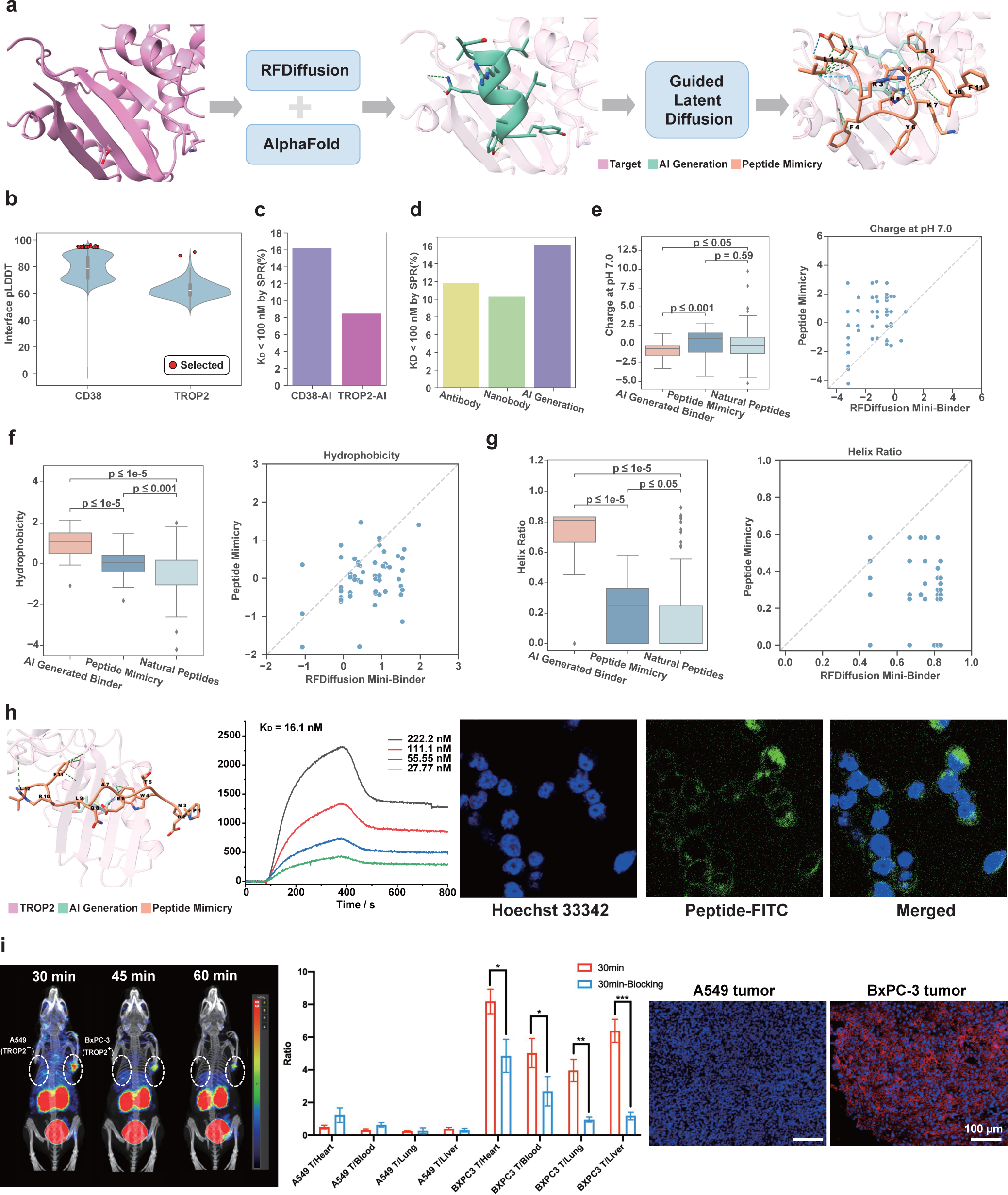
Results of peptide mimicry of AI-generated binding structures. **a**, The overall pipeline for generating peptide mimicries in the absence of reference binders. RFDiffusion is initially used to create mini-binders, followed by AlphaFold Multimer for filtering and structural prediction. The resulting binding structures with high interface pLDDT are then utilized for peptide mimicry design. **b**, Distribution of interface pLDDT scores for the selected binding structures for mimicry. The interface pLDDT is defined as the average pLDDT of residues within 6.0Å C_(3_ distance to the binding partner. **c**, Success rates (K_D_ < 100 nM) of the pipeline across different target proteins. **d**, Success rates categorized by the type of reference binder, with “AI Generation” indicating binders obtained through the pipeline. **e-g** Comparison of physiochemical properties and structural characteristics among molecules from different sources, including isoelectric point, hydrophobicity defined by GRAVY^27^, and helix ratio. The scatter plots profile the change of these properties by mimicking the mini-binders generated by RFDiffusion^47^. The peptide mimicries only include those with K_D_ < 100 nM measured by SPRi. **h**, Affinity and specificity evaluation of the selected TROP2 peptide. SPRi binding curves of the peptides and the TROP2 protein. Confocal imaging of the binding of the selected peptide to BxPC3 cells (TROP2 positive). Scale bar: 20 *µ*m. **i**, Micro PET images obtained at 30, 45 and 60 min post injection of ^68^Ga DOTA-PDMWTEAQLRFL in mice with bilateral A549 and BxPC3 tumors xenografts (n=5). Tumor-to-normal tissue uptake ratio in BxPC3 and A549 tumor-bearing mice. Immunofluorescence staining displaying TROP2 expression in tumor tissues.

As expected, the mini-binders generated by RFDiffusion had large discrepancies from the natural peptides regarding physiochemical properties including isoelectric point, charge at pH 7.0, and hydrophobicity (Supplementary Fig. 9a-c), the last of which might further induce higher aggregation propensity (Supplementary Fig. 9d). RFDiffusion also encouraged the generation of helices, suppressing strands and loops compared to natural peptides (Supplementary Fig. 9e-g). For peptide mimicries based on these mini-binders, the success rates, defined by a K_D_<100 nM, were approximately 16% for CD38 and 8% for TROP2 (Fig. 6c). Further analysis of CD38 success rates, categorized by different reference binders, indicated that using AI-generated mini-binders through this pipeline typically resulted in higher success rates (Fig. 6d). We hypothesized this was because AI-generated mini-binders created smaller and more compact interfaces with the target protein compared to natural binders, such as antibodies or nanobodies (Supplementary Fig. 9h), thereby imposing fewer constraints on peptide mimicries. In-depth analysis revealed that peptide mimicries resembled natural peptides more closely than AI-generated mini-binders in terms of physiochemical properties and secondary structures (Fig. 6e-g). We visualized the structure of one high-affinity binder designed for TROP2, which also demonstrated excellent specificity in cellular assays (Fig. 6h). Confocal imaging confirmed the high and targeted binding of the selected peptide to the membrane surface of BxPC3 cells, which express high levels of TROP2, indicating its high affinity and specificity (Fig. 6h, Supplementary Fig. 10a). The peptide was also selected for *in vivo* validation on tumor mouse models and exhibited potential for therapeutic purposes. Small animal PET/CT imaging demonstrated a 16.18-fold greater uptake of the peptide probe in BxPC3 tumors compared to A549 tumors (6.15 ± 0.9% ID/cc vs. 0.38 ± 0.07% ID/cc) 30 minutes after intravenous injection (Fig. 6i). The BxPC3 tumor side of the dual-tumor mouse model exhibited significant uptake of radioactive signal, whereas the A549 tumor side showed negligible tracer uptake. Immunofluorescence analysis, consistent with PET/CT findings, confirmed high expression of TROP2 in BxPC3 tumors and very low expression in A549 tumors (Fig. 6i). These results demonstrate that ^68^Ga– DOTA – PDMWTEAQLRFL, a novel PET/CT tracer with high affinity and specificity for TROP2, has potential applications in tumor imaging and guiding individualized treatment strategies. Another designed peptide binding to TROP2 also demonstrated good performance, with the similar *in vitro* and *in vivo* experimental results included in Supplementary Fig. 10b-e. We screened the same random peptide library for peptides that can bind to TROP2, and only 9 peptides with an affinity of 10^-8^M were identified. Thus, our model achieved a success rate 90,000 times higher than that of the experimental approach for this target (Methods).

## Discussion

We have developed an innovative geometric generative model for the simultaneous design of peptide sequences and their all-atom structures. Given the complex structure of a target protein and an existing binder, PepMimic is capable of generating peptide mimicries of the binding interface with atomic precision. Unlike models like AlphaFold3^1^ and RosettaFold All-Atom^26^, which focus on structure prediction from sequence, our model focuses on co-designing the sequences and all-atom structures of target-binding peptides. This makes it uniquely suited for the generation of peptide mimicries that reproduce key molecular interactions, especially in complex biological environments.

PepMimic surpassed state-of-the-art protein design models^18, 25, 47^ on high-quality testing protein-peptide complexes from literature^43^. SPRi experiments showed that our designed peptides achieved 8% success rate (defined by dissociation constants, K_D_, below 100 nM) across five targets on average, much higher than that of traditional random library screening. For targets lacking available complex structures, we have also proposed a pipeline that first leverages existing protein binder design models, followed by PepMimic for designing peptide mimicries of the generated interface. The pipeline yielded an average of 14% success rate across two targets, highlighting its potential in challenging scenarios. The motivation for this pipeline stems from the low developability of mini-proteins, which are prone to triggering immune responses due to non-humanized amino acid sequences. Such challenges might be potentially avoided by designing peptide counterparts, which tend to have lower immunogenicity, as the substitute.

PepMimic holds promise for a wide range of biological and therapeutic applications requiring peptides mimicking the function of existing binders, especially in systematically transforming existing antibody or nanobody drugs into peptide analogs with enhanced oral bioavailability and potency. We also envision that the same computational framework could be extended to design antibodies or nanobodies that mimic natural or AI-generated binders. While this study has primarily focused on developing imaging peptides for different tumors, future work will step forward to explore therapeutic peptides for cancer treatment.

## Methods

### Dataset

The training data comprised protein-peptide complexes collected from the Protein Data Bank (PDB)^3^ before December 8th, 2023. We first extracted all dimers from the PDB and retained the complexes with a receptor longer than 30 residues and a peptide between 4 to 25 residues^16^. Next, MMseqs2^15^ was used to derive the sequence identity of the receptors and the peptides, respectively. Complexes with both receptor and peptide sequence identities over 90% were considered duplicates and removed, after which 6,105 unique complexes remained. To ensure a cross-target generalization test, we leveraged 93 complexes from the Large Non-Redundant (LNR) dataset curated in previous literature^16^ as the test set. All complexes were clustered by receptor sequence identity with a threshold of over 40%, and those sharing clusters with test complexes were excluded to prevent data leakage. The remaining data were randomly split into a training set of 4,157 complexes and a validation set of 114 complexes based on the clustering results.

An unsupervised dataset was also constructed for pre-training, comprising 70,645 peptide-like protein fragments from monomers. Specifically, we extracted all single chains from the PDB, and removed duplicated chains with a sequence identity threshold over 90%. For each chain, the fragments satisfying the following criteria were selected. (1) **Length**: The fragment should consist of 4 to 25 residues. (2) **Balanced composition**: No single amino acid should constitute more than 25% of the fragment; Hydrophobic amino acids should comprise less than 45% of a given fragment, with charged amino acids accounting for 25% to 45%. (3) **Isolated Stability**: Instability^7^ should be below 40; Considering surrounding amino acids as interaction partners, the fragment should have a buried surface area (BSA)^4^ above 400Å^2^, with a relative BSA above 20%. FreeSASA^13^ was used to calculate the surface area of the fragments. During training, residues with C_(3_ distances to peptide residues below 10Å were designated as the binding site.

### Model Architecture

PepMimic integrates three core components: an autoencoder (*ε_ϕ_, D_ξ_*) to derive a low-dimensional latent space, a diffusion model *ϵ_θ_* defined in the latent space, and a latent interface encoder ***f**ψ* used for guidance to mimic the provided binding interfaces (Fig. 1b-c). PepMimic first leverages an autoencoder^10^ to establish a reversible mapping from the all-atom geometries of peptides to a compressed and standardized latent space (Supplementary Notes 1.3.2). The coordinates in the latent space further endure a target-specific invertible affine transformation *F* derived from the covariance matrix 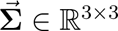 of the binding site *𝒢_b_* via the Cholesky decomposition^6^:

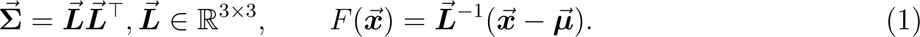

This affine transformation *F* maps all data distributions to a similar scale, and thus enhances the stability in training and inference of the diffusion model, preventing generated coordinates from expanding to infinity or shrinking to the origin. Subsequently, PepMimic implements a diffusion model^8^ within this latent space to generate meaningful latent point clouds from a standard Gaussian distribution. Compared to RFDiffusion^17^ which only generates backbone coordinates with diffusion, the autoencoder and the affine transformation in PepMimic defined a well-formed joint latent space for sequences and all-atom structures, facilitating the learning of the diffusion model for co-design. Building upon the latent representations produced by the autoencoder, a latent interface encoder was further trained to distinguish similar interfaces from dissimilar ones, which are constructed by the downsampling strategy (Fig. 1d, Supplementary Notes 1.2.1). The distances of the representations of different interfaces provided by the latent interface encoder reflect their similarities. Given a target protein and a specific binder (e.g. an antibody) to mimic, we initially extracted their interface to derive the reference latent point cloud, and then obtain a vector-form representation 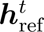 via the latent interface encoder. Then, for each state of the latent point clouds during generation, its vector-form representation is similarly derived. The distance on these two latent representations is used to guide the generative diffusion process, minimizing the distance via gradient guidance (Supplementary Notes 1.6.2):

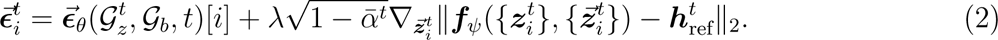

Consequently, the generated peptide gradually aligns with the desired interface in the latent space. By decoding the latent point clouds back to all-atom geometries via the autoencoder, we obtain a peptide binder that mimics the interface between the target and the given binder (Fig. 1a,c). The autoencoder and the diffusion model were parameterized with three-layer dynamic multi-channel equivariant graph neural networks^11^ (Supplementary Notes 1.3.5), which was designed for modeling all-atom geometries. The latent interface encoder was parameterized with a three-layer equivariant transformer (Supplementary Notes 1.3.6) to better capture the overall shapes of the latent point clouds. More technical details on hyperparameters could be found in the Supplementary Notes 4.

### Training Regimen

The all-atom autoencoder was trained on the dataset merging the 70K peptide-like protein fragments and the 4k protein-peptide complexes. The objective *L*_AE_ includes *L*_KL_, the KL divergence between the latent embedding and the prior distribution, as well as *L*_rec_, the reconstruction loss of the amino acid types and the all-atom geometries. The prior distributions for amino-acid-level latent embeddings are *N*(0, I) and 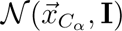, respectively, where 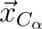 is the coordinate of its alpha carbon. The reconstruction loss includes supervision on the atom-level coordinates, bond lengths, and side-chain torsional angles. The latent diffusion model was first pre-trained on the protein fragments and then fine-tuned on the protein-peptide complexes, with denoising objectives^8^ on both the invariant and the equivariant latent embeddings of amino acids on the peptides. For training the latent interface encoder, we resorted to down-sampling interfaces from the protein fragments and protein-peptide complexes to construct a corresponding dataset (Fig. 1d). Initially, amino acids on the peptides were randomly split into two non-overlapping sets, A and B. Set A is down-sampled twice to create a positive pair with overlapping amino acids, while set B is down-sampled once to form a negative sample without any overlapping amino acids with the former two samples. The latent interface encoder was then trained to minimize the L_2_ distances between the representations of positive pairs and maximize those between negative pairs (Fig. 1d). Details of the training objectives for the autoencoder, diffusion model, and the interface encoder are provided in Supplementary Notes 1.4. Both the autoencoder and the latent interface encoder were trained for 100 epochs using AdamW optimizer with an initial learning rate of 1 10^-4^ and a scheduler that decayed the learning rate by 20% after five epochs without improvement on the validation set. The pre-training and the fine-tuning phase of the latent diffusion model last for 100 epochs with initial learning rates of 1 × 10^-4^ and 5 × 10^-5^, respectively.

### Inference Regimen

Given the binding site as the condition, we first moved its center of mass to the origin of the coordinate system, and sampled a random latent point cloud from the standard Gaussian distribution. The latent diffusion model then denoised the latent point clouds for *T* (= 100) steps to produce a meaningful result, which was subsequently decoded by the autoencoder into amino acid types and all-atom structures. For mimicking the interaction of a specific binder to the target protein, we first extracted their interface, defined by amino acids on the binding site within 10Å *C_β_* distances to the binder, and residues on the binder within 6Å atomic distances to the binding site. This interface was sequentially fed into the autoencoder and the latent interface encoder to derive a vector-form representation h_ref_ ∈ ℝ*^d^*. During generation, the latent point cloud at time step *t* is also projected into an interface representation h*_t_* ∈ ℝ*^d^* via the latent interface encoder. The distance between h_ref_ and h*_t_* was calculated and its gradients with respect to the invariant and the equivariant latent embeddings were used to adjust the latent point cloud. This process aligns the latent point clouds with the reference interface in the latent space, which produces a peptide mimicry after decoding back to all-atom geometries by the autoencoder.

### Evaluation Metrics

We calculated the Amino Acid Recovery (AAR), Root Mean Square Deviation (RMSD), DockQ^2^, and Rosetta Interface Energy (Δ*G*)^1^ for *In silico* evaluations on target-specific peptide design. AAR, RMSD, and DockQ were computed by comparing the generated peptide with the reference peptide binder. AAR assesses the similarity between the generated sequence and the reference sequence by calculating the percentage of matching amino acids. As the sequences do not need to align perfectly from the start (N-terminus), we slide the generated sequence along the reference sequence to obtain the highest AAR score. RMSD measures the deviation of the generated C_*α*_ coordinates from the reference coordinates. DockQ^2^ evaluates the all-atom similarity between the interfaces formed by the target protein with the generated peptide and the reference peptide. For each target protein, we generated 40 candidates and selected the best scores for each metric to assess how well the generated distribution recovered the reference distribution. Rosetta interface energy was calculated using the default REF2015^1^ score function, with the generated complexes undergoing the FastRelax^12^ protocol in advance.

### Ranking Strategy for SPRi Experiments

Rosetta interface energy^1^, FoldX energy^14^, Interface Hit (defined as before), and pLDDT score from AlphaFold Multimer^5, 9^ were employed to rank and select the most promising candidates for further experiments. Initially, basic thresholds were set to filter out candidates with Rosetta interface energy above -10.0, FoldX energy above -5.0, or Interface Hit below 2.0. Candidates failing to meet these thresholds were considered either physically invalid or ineffective at mimicking the reference interface. The ranks from Rosetta and FoldX were averaged to select the top 1,000 candidates for further evaluation with AlphaFold Multimer, due to practical consideration of computational cost. Candidates with pLDDT scores above 70.0 for the peptide, along with top 500 candidates from Rosetta and FoldX rankings, respectively. The selected peptides were hierarchically clustered using a sequence alignment score threshold of 0.4. The sequence alignment score was calculated via dynamic programming, with global alignment mode and scoring parameters set as follows: match = 1, mismatch = 0, open gap = -2, and extend gap = -1. Scores were then normalized by the square root of the product of the lengths of the two sequences. Then each candidate was assigned a priority, with the highest priority (-1) given to those with pLDDT scores above 70.0. Other candidates were prioritized based on the smaller ranking between Rosetta and FoldX. In each cluster, only the candidate with the highest priority was retained, while others were discarded. The remaining candidates were re-ranked based on priority, and the top K candidates were chosen for SPRi experiments. For targets with available reference binders, K was set to 384. For pipeline-based experiments, K was set to 94 for TROP2 and 290 for CD38, totaling 384. Additional implementation details of Rosetta, FoldX, and Alphafold Multimer can be found in Supplementary Notes 4.

### SPRi Experiments

SPRi analysis was conducted using a Plexera PlexArray HT system (Plexera LLC, Bothell, WA). Peptides were immobilized on the gold chip surface, and nonspecific binding was minimized by treatment with 5% nonfat milk. The assays utilized protein concentrations ranging from 1.25 *µ*g/mL to 20 *µ*g/mL in PBST buffer, introduced to the chip at a flow rate of 2 *µ* L/min. Binding was quantified in response units, with association and dissociation times of 300 seconds each. Chip regeneration was performed using 0.5%(v/v) H3PO4 in deionized water. The affinity constant (KD) was calculated using the PlexArray HT system.

### *In vitro* fluorescent imaging

The H1975, MDA-MB-231, BxPC3, MCF-7, and 293T cell lines were cultured in RPMI 1640 medium or DMEM/high glucose supplemented with 10% FBS. The cells were then digested, and approximately 1 10^5^ cells/mL were seeded into culture dishes and incubated overnight to allow for the cell adherence. FITC-labeled peptide was prepared in PBS at a concentration of 20 *µ*M. The cells were incubated with 100 *µ*L of FITC-labeled peptide solution containing Hoechst 33342 (1 mM) in the dark for 30 minutes at 4°C. Finally, the cells were washed three times with cold PBS. Fluorescence imaging was performed using a laser confocal microscope.

### *In vivo* imaging of tumor

NOD/SCID mice were subcutaneously injected with 100 *µ*L of saline containing various cancer cells into the upper right flank. When the tumor volume reached approximately 150 mm^3^, each mouse received a tail vein injection of 200 *µ*L (50 *µ*M) of PBS solution containing the ICG-labeled peptide, while the control group was injected with the same concentration of pure ICG in PBS solution. 30 minutes post-injection, the mice were anesthetized with 4% chloral hydrate and placed in an IVIS Imaging System for detection. Additionally, the peptide was conjugated with the bifunctional chelator 1,4,7,10-tetraazacyclododecane-1,4,7,10-tetraacetic acid (DOTA), as previously described. For radionuclide labeling, ^68^Ga was added to a NaOAc solution (pH 5.5) containing the peptide-DOTA conjugate and reacted at 90°C for 10 minutes. The ^68^Ga-labeled peptide was then purified using a C18 column. Tumor models were intravenously injected with the ^68^Ga-labeled peptide (8-10 MBq per mouse, n=3) via the tail vein, and PET imaging was performed using a microPET/CT scanner (Mediso). In cold-probe blocking experiments, a non-labeled peptide (50 *µ*g) was co-injected with the tracer. Regions of interest (ROIs) over the tumor sites were delineated to calculate the percentage of the injected dose per cubic centimeter (%ID/cc) based on the average signal intensity within these ROIs. For biodistribution studies, tumors and other tissues were harvested and analyzed. The radioactivity in each sample was measured using a I counter. Tumors were fixed in 4% paraformaldehyde and sectioned. The sections were washed with TBST solution and incubated overnight with anti-human PD-L1 antibodies (1:500, Abcam). This was followed by incubation with Cy3-labeled goat anti-mouse IgG (1:2000, Servicebio) as the secondary antibody for 1 hour and with DAPI for 2 minutes. The sections were then observed using a fluorescence microscope.

### Peptide chemical synthesis library

In order to compare with AI-designed peptides, we constructed a random peptide library containing peptides targeting both PD-L1 and TROP2. Using Tenta Gel resin as the solid-phase support, we applied the Fmoc solid-phase synthesis method to create peptides with the sequence X_1_X_2_X_3_X_4_X_5_X_6_X_7_X_8_X_9_X_10_M. The library was synthesized using a “mix-and-split” strategy: at each step, the products were evenly divided, different amino acids were randomly added, thoroughly mixed, and divided again. This cycle continued until the final peptide sequence was completed. The synthesis was conducted using a peptide synthesizer, with each coupling reaction including the following steps: (1) Deprotection: Removal of the Fmoc group using 20% piperidine; (2) Washing: Cleaning with N, N-dimethylformamide (DMF); (3) Coupling: Adding Fmoc-protected amino acids dissolved in a 4% N-methyl morpholine DMF solution, combined with an equivalent amount of 2-(1H-Benzotriazol-1-yl)-1,1,3,3-tetramethyluronium Hexafluoro-phosphate (HBTU). After coupling, deprotection and washing were repeated, followed by thorough resin mixing. Final side-chain deprotection was achieved with 95% trifluoroacetic acid (TFA), and the resin was washed with methanol and vacuum-dried. The peptide library was incubated with biotin-labeled PD-L1 and TROP2 protein and streptavidin magnetic beads were added. Finally, the positive peptides were separated by magnetic field and then identified to obtain the peptide sequences.

## Data Availability

All the training data, the test set, the unsupervised dataset, and detailed splits, can be found at Zenodo. The complexes used for mimicry are extracted from PDB (https://www.rcsb.org/), the PDB IDs of which are listed in Supplementary Table 1-5.

## Code Availability

Codes for running our peptide mimicry algorithm are accessible at https://github.com/ kxz18/PepMimic.

## Author contributions

X.K. and J.M. conceived and designed the generative model with contributions from R.J., R.G. and J.G. X.P., R.G., J.G., and Y.J. processed the data. X.P., R.G., and J.G. conducted computational experiments. Z.W. conducted the wet-lab experiments. X.K., M.Z., Z.W., and J.M. analyzed the results. X.K., R.J., Z.W., and J.M. wrote the manuscript. The study was supervised by W.H., Y.L., and J.M.

