## supplementary materials for "Peptide Design through Binding Interface Mimicry"

**2 Baseline Implementations** **17**
**3 Normalized Rosetta Interface Energy** **18**
**4 Implementation Details** **18**

#### List of Algorithms

1 Algorithm for Downsampling Interfaces in Latent Space . . . . . . . . . . 3
2 Algorithm for Acquring Positive and Negative Interface Pairs . . . . . . . . . . 4
3 Adaptive Multi-Channel Equivariant Graph Neural Network . . . . . . . . . . 10
4 Geometric Equivariant Transformer . . . . . . . . . . 11
5 Training Algorithm of All-Atom Variational AutoEncoder . . . . . . . . . . 13
6 Training Algorithm of Latent Diffusion Model . . . . . . . . . . 14
7 Training Algorithm of Latent Interface Encoder . . . . . . . . . . 15
8 Sampling Algorithm of PepMimic . . . . . . . . . . 16

9 Guided Sampling Algorithm of PepMimic . . . . . . . . . . . . . . . . . . . . . . . . . . . . . . . . . . . . . . . . . 17

### 1 Supplementary Methods

#### 1.1 Notations

We detail the algorithms of each module in the subsequent sections, with the following key notations. We denote the training dataset as $\mathcal{S}$. $\mathcal{E}_\phi, \mathcal{D}_\xi, \boldsymbol{\epsilon}_\theta, \mathcal{E}_\mathcal{I}$ represent the learnable all-atom encoder, decoder, latent diffusion model and latent interface encoder, parameterized by $\phi, \xi, \theta$ and $\mathcal{I}$ respectively. We use $\mathcal{G}_p$, $\mathcal{G}_b$, $\tilde{\mathcal{G}}_p$, and $\mathcal{G}_z$ to represent the geometric graphs corresponding to the data-space peptide, the context environment (i.e. the binding site), the masked peptide, and the latent graph of the peptide. We denote $\boldsymbol{H}, \vec{\boldsymbol{X}}, \vec{\mathbf{X}}$ as the matrix of invariant scalers, equivariant vectors, and a collection of vector matrices. $\vec{\boldsymbol{u}}_i^t = (\boldsymbol{z}_i^t, \vec{\boldsymbol{z}}_i^t)$ is the latent representation of node $i$, timestep $t$ with both scaler and vector features. For the details of model architectures, we use $\varphi$ (with proper subscripts) as Multi-Layer Perceptions (MLPs), and $\mathrm{RBF}, \mathrm{Softmax}, \mathrm{LayerNorm}, \mathrm{CrossEntropy}, \mathrm{MSE}$ denote the Radial Basis Functions, the Softmax activation, the Layer Normalization, the Cross Entropy loss function, and the Mean Square Error. $\|\cdot\|_2$ and $|\cdot|$ are the L2-norm of a vector and the cardinal of a set. For distributions, we use $\mathbf{U}(a, b)$ to denote uniform distribution between $a$ and $b$, as well as $\mathcal{N}(\mu, \sigma^2)$ to denote Gaussian distribution with expectation $\mu$ and variance $\sigma^2$.

#### 1.2 Data Processing

##### 1.2.1 Downsampling Latent Interface

The downsampling strategy are used in data construction for evaluation on the reliability of the proposed interface hit, as well as the training and testing of the latent interface encoder. It keeps 50% to 100% amino acids in the original graph of the interface, represented as a latent point cloud by the all-atom encoder $\mathcal{E}_\phi$, to produce a subgraph, with detailed description provided in Algorithm 1.

**Algorithm 1** Algorithm for Downsampling Interfaces in Latent Space

```
def DownSampling(𝒢):
#   Determine a Preservation Ratio between 50% and 100%
    R ~ U(0.5, 1)
    𝒢_sub ← ∅
    for i in 1, 2, ··· , |𝒢| do
        r ~ U(0, 1)
        if r ≤ R then
            𝒢_sub ← 𝒢_sub ∪ {(z_i, z⃗_i)}
        end if
    end for
    return 𝒢_sub
```

##### 1.2.2 Construction of Test Interface Pairs via Partitioning and Downsampling

To train and evaluate the latent interface encoder for further guided sampling, we adopt a contrastive strategy by encouraging the encoder to yield difference representations of two non-overlapping point clouds. To obtain such pairs, we resort to partitioning and downsampling strategies on the interfaces. Specifically, given a latent geometric graph, we first randomly partition the amino acids, represented as a latent point cloud, into two distinct sets A and B, with equal probability. Using a subgraph downsampled (Supplementary Notes 1.2.1) from A as the reference, we sample another subgraph from the same set A as the positive graph, while the complementary set B is used to create the negative graph. The sampled subgraphs are then perturbed to timestep $t$ to match with the states in the diffusion generative process. Detailed algorithms for constructing the interface pairs are provided in Algorithm 2.

**Algorithm 2** Algorithm for Acquring Positive and Negative Interface Pairs

**def** PairSampling($\mathcal{G}$):<br>
  $\mathcal{G}_+, \mathcal{G}_- \leftarrow \emptyset, \emptyset$<br>
  # *Split the Original Graph with Equal Probability*<br>
  **for** $i$ in $1, 2, \cdots, |\mathcal{G}|$ **do**<br>
    $r \sim \mathbf{U}(0, 1)$<br>
    **if** $r \leq 0.5$ **then**<br>
      $\mathcal{G}_+ \leftarrow \mathcal{G}_+ \cup \{(\boldsymbol{z}_i, \vec{\boldsymbol{z}}_i)\}$<br>
    **else**<br>
      $\mathcal{G}_- \leftarrow \mathcal{G}_- \cup \{(\boldsymbol{z}_i, \vec{\boldsymbol{z}}_i)\}$<br>
    **end if**<br>
  **end for**<br>
  $\mathcal{G}_{\text{ref}} \leftarrow$ DownSampling($\mathcal{G}_+$)<br>
  $\mathcal{G}_+ \leftarrow$ DownSampling($\mathcal{G}_+$)<br>
  $\mathcal{G}_- \leftarrow$ DownSampling($\mathcal{G}_-$)<br>
  **return** $\mathcal{G}_{\text{ref}}, \mathcal{G}_+, \mathcal{G}_-$

#### 1.3 Model Architecture

The overall workflow of our PepMimic consists of four modules: (1) An all-atom variational autoencoder that defines the joint latent space for sequences and all-atom structures conditioned on the all-atom context of the binding site (Supplementary Notes 1.3.1); (2) An affine transformation derived from the binding site to project the 3D geometry into a standard space approximating standard Gaussian distribution (Supplementary Notes 1.3.2); (3) A latent diffusion model trained on the standard latent space (Supplementary Notes 1.3.3). (4) A latent interface encoder trained to distinguish similar and dissimilar interfaces in the latent space (Supplementary Notes 1.3.4).

##### *1.3.1 All-Atom Variational AutoEncoder*

The all-atom autoencoder consists of an encoder $\mathcal{E}_\phi$ that encodes the peptide $\mathcal{G}_p$ in the presence of the binding site $\mathcal{G}_b$ into a latent point cloud $\mathcal{G}_z$, and a decoder $\mathcal{D}_\xi$ that reconstructs the peptide from the latent state to obtain $\mathcal{G}'_p = \{(x'_i, \vec{\boldsymbol{X}}'_i)\}$. To encourage $\mathcal{E}_\phi$ to learn contextual representations of residues, we corrupt 25% of the residues in $\mathcal{G}_p$ with a [MASK] type to obtain $\tilde{\mathcal{G}}_p$ as the input:

$$\mathcal{G}_z = \mathcal{E}_\phi(\tilde{\mathcal{G}}_p, \mathcal{G}_b), \qquad \mathcal{G}'_p = \mathcal{D}_\xi(\mathcal{G}_z, \mathcal{G}_b), \tag{3}$$

where $\mathcal{G}_z = \{(\boldsymbol{z}_i, \vec{\boldsymbol{z}}_i) | i \in \mathcal{G}_p\}$ contains the latent states $\boldsymbol{z}_i \in \mathbb{R}^h$ ($h = 8$ in this paper) and $\vec{\boldsymbol{z}}_i \in \mathbb{R}^3$ sampled from the encoded distribution $\mathcal{N}(\boldsymbol{z}_i; \boldsymbol{\mu}_i, \boldsymbol{\sigma}_i)$ and $\mathcal{N}(\vec{\boldsymbol{z}}_i; \vec{\boldsymbol{\mu}}_i, \vec{\boldsymbol{\sigma}}_i)$ using the reparameterization trick[13]. We borrow the adaptive multi-channel equivariant graph neural network (Supplementary Notes 1.3.5) in dyMEAN[15] for both $\mathcal{E}_\phi$ and $\mathcal{D}_\xi$ to capture the all-atom geometry. In the decoder $\mathcal{D}_\xi$, we factorize the joint distribution of sequences and all-atom structures as follows:

$$p_\xi(x'_i, \vec{\boldsymbol{X}}'_i | \mathcal{G}_z, \mathcal{G}_b) = p_{\xi_1}(x'_i | \mathcal{G}_z, \mathcal{G}_b) p_{\xi_2}(\vec{\boldsymbol{X}}'_i | x'_i, \mathcal{G}_z, \mathcal{G}_b), \tag{4}$$

where the sequence is first decoded from the latent states and then the all-atom geometry, initialized with replications of $\vec{z}_i$, is reconstructed. The training objectives involves the reconstruction loss $\mathcal{L}_{recon}$ and the KL divergence $\mathcal{L}_{KL}$ to constrain the latent space (Supplementary Notes 1.4.1). And the training procedure is depicted in Supplementary Notes 1.5.1.

###### 1.3.2 Target-Specific Affine Transformation

With the latent space given by the autoencoder, we further exploit a standard space obtained from target-specific affine transformations, which enhances the transferability of diffusions on disparate binding sites. Most peptides fold into complementary shape upon binding on the receptor[16,20]. Thus, the target distribution is inherently characterized by the shape of the binding site. Given the wide disparity in binding geometries, directly implementing diffusion in the data space yields minimal transferability among different binding sites. To address this deficiency, we propose to implement the diffusion process on a shared standard space converted via an affine transformation derived from the binding site. Formally, denoting the $\text{C}_\alpha$ coordinates of the residues in a given binding site $\mathcal{G}_b$ as $\vec{\boldsymbol{R}} \in \mathbb{R}^{3\times|\mathcal{G}_b|}$, we can derive their center $\vec{\boldsymbol{\mu}} = \mathbb{E}[\vec{\boldsymbol{R}}] \in \mathbb{R}^3$ and covariance $\vec{\boldsymbol{\Sigma}} = \text{Cov}(\vec{\boldsymbol{R}}, \vec{\boldsymbol{R}}) \in \mathbb{R}^{3\times 3}$, so that these coordinates can be regarded as sampled from the distribution $\mathcal{N}(\vec{\boldsymbol{\mu}}, \vec{\boldsymbol{\Sigma}})$. We then calculate the Cholesky decomposition[7] of $\vec{\boldsymbol{\Sigma}}$:

$$\vec{\boldsymbol{\Sigma}} = \vec{\boldsymbol{L}}\vec{\boldsymbol{L}}^\top, \vec{\boldsymbol{L}} \in \mathbb{R}^{3\times 3}, \tag{5}$$

where $\vec{\boldsymbol{L}}$ is a lower triangular matrix. $\vec{\boldsymbol{L}}$ is unique[7] and invertible since the covariance matrix is a real-valued symmetric positive-definite matrix[1]. Then we can define the affine transformation $F : \mathbb{R}^3 \to \mathbb{R}^3$, which enables the projection of the geometry into the standard space approximating standard Gaussian $F(\vec{\boldsymbol{R}}) \sim \mathcal{N}(\boldsymbol{0}, \boldsymbol{I})$. Further, we can easily obtain the inverse of $F$ as:

$$F(\vec{\boldsymbol{x}}) = \vec{\boldsymbol{L}}^{-1}(\vec{\boldsymbol{x}} - \vec{\boldsymbol{\mu}}), \qquad F^{-1}(\vec{\boldsymbol{x}}) = \vec{\boldsymbol{L}}\vec{\boldsymbol{x}} + \vec{\boldsymbol{\mu}}. \tag{6}$$

With the above definitions, for each given binding site $\mathcal{G}_b$, we transform the geometry via the derived $F$ to obtain the standard space, where the diffusion model is implemented, and recover the original geometry with $F^{-1}$ (Fig. 1a) after generation. Notably, we have the following proposition to ensure that the equivariance is maintained under the proposed affine transformation with scalarization-based equivariant GNNs[4,8]:

**Proposition 1.1.** *Denote the invariant and equivariant outputs from a scalarization-based E(3)-equivariant GNN as $f(\{\boldsymbol{h}_i, \vec{\boldsymbol{x}}_i\})$ and $\vec{f}(\{\boldsymbol{h}_i, \vec{\boldsymbol{x}}_i\})$, respectively. With the definition of $F$ in Eq. 6, $\forall g \in E(3)$, we have $f(\{\boldsymbol{h}_i, F(\vec{\boldsymbol{x}}_i)\}) = f(\{\boldsymbol{h}_i, F_g(g\cdot\vec{\boldsymbol{x}}_i)\})$ and $g \cdot F^{-1}(\vec{f}(\{\boldsymbol{h}_i, F(\vec{\boldsymbol{x}}_i)\})) = F_g^{-1}(\vec{f}(\{\boldsymbol{h}_i, F_g(g\cdot\vec{\boldsymbol{x}}_i)\}))$, where $F_g$ is derived on the coordinates transformed by $g$. Namely, the E(3)-equivariance is preserved if we implement the GNN on the standard space and recover the original geometry from the outputs.*

This is vital since it indicates the Markov kernel is E(3)-equivariant, and thus ensures the E(3)-invariance of the probability density in the diffusion process[22]. Note that our variational autoencoder and latent diffusion model are already designed to be equivariant even without the proposed affine transformation here. The purpose of defining such component is to encourage better generalization of the diffusion processes. Indeed, it is nontrivial to analyze whether such an

[1] The binding site has at least 3 non-overlapping nodes, namely $\text{rank}(\vec{\boldsymbol{R}}) = 3$, thus we can ignore the corner case of semi-positive definite matrices.

implementation will break the equivariance of our workflow. Luckily, Proposition 1.1 manages to prove that scalarization-based equivariant networks[8] are seamlessly compatible with such affine transformation, which naturally preserves equivariance without any requirements of adaption.

Here we provide the proof of Proposition 1.1 as follows. For simplicity, given $F$ derived from a set of coordinates $\{\vec{\boldsymbol{x}}_i\}$ following Eq. 6, we use $F_g$ to indicate the affine transformation derived from $\{g \cdot \vec{\boldsymbol{x}}_i\}$, where $g \in E(3)$. Additionally, we keep the terminology "standard space" for describing the space after the data-specific affine transformation $F$. We begin by proving a key lemma, that is, the E(3)-invariance of distances between two nodes in the standard space converted by $F$:

**Lemma 1.2.** *Given two nodes $i$ and $j$ in the geometric graph $\mathcal{G}$, denoting their coordinates as $\vec{\boldsymbol{x}}_i$ and $\vec{\boldsymbol{x}}_j$, their distance in the standard space is E(3)-invariant. Namely, $\forall g \in E(3), \|F(\vec{\boldsymbol{x}}_i) - F(\vec{\boldsymbol{x}}_j)\| = \|F_g(g \cdot \vec{\boldsymbol{x}}_i) - F_g(g \cdot \vec{\boldsymbol{x}}_j)\|$.*

*Proof.* $\forall g \in E(3)$, $g$ can be instantiated as an orthogonal matrix $\boldsymbol{Q} \in O(3)$ (including rotation and reflection), and a translation vector $\vec{\boldsymbol{t}} \in \mathbb{R}^3$. Denoting all coordinates in the geometric graph as $\vec{\boldsymbol{X}} \in \mathbb{R}^{3\times|\mathcal{G}|}$, and the number of nodes in $\mathcal{G}$ as $n$, we can derive the E(3)-equivariance of the expectation of the coordinates:

$$\begin{aligned}
\mathbb{E}[g \cdot \vec{\boldsymbol{X}}] &= \frac{1}{n}\sum_i g \cdot \vec{\boldsymbol{x}}_i = \frac{1}{n}\sum_i (\boldsymbol{Q}\vec{\boldsymbol{x}}_i + \vec{\boldsymbol{t}}) && (7)\\
&= \frac{1}{n}\boldsymbol{Q}(\sum_i \vec{\boldsymbol{x}}_i) + \vec{\boldsymbol{t}} = \boldsymbol{Q}\mathbb{E}[\vec{\boldsymbol{X}}] + \vec{\boldsymbol{t}} && (8)\\
&= g \cdot \mathbb{E}[\vec{\boldsymbol{X}}]. && (9)
\end{aligned}$$

With the E(3)-equivariance of the expectation, it is easy to derive the following equation on the covariance matrix:

$$\begin{aligned}
\mathrm{Cov}(g \cdot \vec{\boldsymbol{X}}, g \cdot \vec{\boldsymbol{X}}) &= \frac{1}{n-1}(g \cdot \vec{\boldsymbol{X}} - \mathbb{E}[g \cdot \vec{\boldsymbol{X}}])(g \cdot \vec{\boldsymbol{X}} - \mathbb{E}[g \cdot \vec{\boldsymbol{X}}])^\top && (10)\\
&= \frac{1}{n-1}(g \cdot \vec{\boldsymbol{X}} - g \cdot \mathbb{E}[\vec{\boldsymbol{X}}])(g \cdot \vec{\boldsymbol{X}} - g \cdot \mathbb{E}[\vec{\boldsymbol{X}}])^\top && (11)\\
&= \frac{1}{n-1}(\boldsymbol{Q}(\vec{\boldsymbol{X}} - \mathbb{E}[\vec{\boldsymbol{X}}]))(\boldsymbol{Q}(\vec{\boldsymbol{X}} - \mathbb{E}[\vec{\boldsymbol{X}}]))^\top && (12)\\
&= \frac{1}{n-1}\boldsymbol{Q}(\vec{\boldsymbol{X}} - \mathbb{E}[\vec{\boldsymbol{X}}])(\vec{\boldsymbol{X}} - \mathbb{E}[\vec{\boldsymbol{X}}])^\top\boldsymbol{Q}^\top && (13)\\
&= \boldsymbol{Q}\mathrm{Cov}(\vec{\boldsymbol{X}}, \vec{\boldsymbol{X}})\boldsymbol{Q}^\top. && (14)
\end{aligned}$$

Based on the Cholesky decomposition used in the derivation of $F$, we denote $\mathrm{Cov}(\vec{\boldsymbol{X}}, \vec{\boldsymbol{X}}) = \vec{\boldsymbol{L}}\vec{\boldsymbol{L}}^\top$ and $\mathrm{Cov}(g \cdot \vec{\boldsymbol{X}}, g \cdot \vec{\boldsymbol{X}}) = \vec{\boldsymbol{L}}_g\vec{\boldsymbol{L}}_g^\top$, with which we can immediately derive:

$$\vec{\boldsymbol{L}}_g\vec{\boldsymbol{L}}_g^\top = \boldsymbol{Q}\vec{\boldsymbol{L}}\vec{\boldsymbol{L}}^\top\boldsymbol{Q}^\top. \tag{15}$$

Considering $\boldsymbol{Q}^{-1} = \boldsymbol{Q}^\top$, we further have the following equation:

$$\vec{\boldsymbol{L}}_g^{-\top}\vec{\boldsymbol{L}}_g^{-1} = \boldsymbol{Q}\vec{\boldsymbol{L}}^{-\top}\vec{\boldsymbol{L}}^{-1}\boldsymbol{Q}^\top. \tag{16}$$

Given that $F(\vec{\boldsymbol{x}}) = \vec{\boldsymbol{L}}^{-1}(\vec{\boldsymbol{x}} - \mathbb{E}[\vec{\boldsymbol{X}}])$, we are now ready to prove Lemma 1.2 as follows:

$$\begin{aligned}
\|F_g(g\cdot\vec{\boldsymbol{x}}_i) - F_g(g\cdot\vec{\boldsymbol{x}}_j)\| &= \|\vec{\boldsymbol{L}}_g^{-1}(g\cdot\vec{\boldsymbol{x}}_i - \mathbb{E}[g\cdot\vec{\boldsymbol{X}}]) - \vec{\boldsymbol{L}}_g^{-1}(g\cdot\vec{\boldsymbol{x}}_j - \mathbb{E}[g\cdot\vec{\boldsymbol{X}}])\| && (17)\\
&= \|\vec{\boldsymbol{L}}_g^{-1}(g\cdot\vec{\boldsymbol{x}}_i - g\cdot\vec{\boldsymbol{x}}_j)\| = \|\vec{\boldsymbol{L}}_g^{-1}\boldsymbol{Q}(\vec{\boldsymbol{x}}_i - \vec{\boldsymbol{x}}_j)\| && (18)\\
&= \sqrt{(\vec{\boldsymbol{L}}_g^{-1}\boldsymbol{Q}(\vec{\boldsymbol{x}}_i - \vec{\boldsymbol{x}}_j))^\top(\vec{\boldsymbol{L}}_g^{-1}\boldsymbol{Q}(\vec{\boldsymbol{x}}_i - \vec{\boldsymbol{x}}_j))} && (19)\\
&= \sqrt{(\vec{\boldsymbol{x}}_i - \vec{\boldsymbol{x}}_j)^\top\boldsymbol{Q}^\top\vec{\boldsymbol{L}}_g^{-\top}\vec{\boldsymbol{L}}_g^{-1}\boldsymbol{Q}(\vec{\boldsymbol{x}}_i - \vec{\boldsymbol{x}}_j)} && (20)\\
&= \sqrt{(\vec{\boldsymbol{x}}_i - \vec{\boldsymbol{x}}_j)^\top\vec{\boldsymbol{L}}^{-\top}\vec{\boldsymbol{L}}^{-1}(\vec{\boldsymbol{x}}_i - \vec{\boldsymbol{x}}_j)} && (21)\\
&= \sqrt{(\vec{\boldsymbol{L}}^{-1}(\vec{\boldsymbol{x}}_i - \vec{\boldsymbol{x}}_j))^\top(\vec{\boldsymbol{L}}^{-1}(\vec{\boldsymbol{x}}_i - \vec{\boldsymbol{x}}_j))} = \|\vec{\boldsymbol{L}}^{-1}(\vec{\boldsymbol{x}}_i - \vec{\boldsymbol{x}}_j)\| && (22)\\
&= \|\vec{\boldsymbol{L}}^{-1}(\vec{\boldsymbol{x}}_i - \mathbb{E}[\vec{\boldsymbol{X}}]) - \vec{\boldsymbol{L}}^{-1}(\vec{\boldsymbol{x}}_j - \mathbb{E}[\vec{\boldsymbol{X}}])\| = \|F(\vec{\boldsymbol{x}}_i) - F(\vec{\boldsymbol{x}}_j)\| && (23)
\end{aligned}$$

□

With Lemma 1.2, we are able to give the proof of Proposition 1.1 as follows.

*Proof.* A first observation is that to prove Proposition 1.1, we only need to prove the equivariance in the 1-layer case, since the multi-layer case can be decomposed into the 1-layer case by inserting $I = F \circ F^{-1}$ between layers, where $I$ is the identical mapping. Generally, each layer in scalarization-based E(3)-equivariant GNN has the following paradigm:

$$\begin{aligned}
\boldsymbol{m}_{ij} &= \phi_m(\boldsymbol{h}_i, \boldsymbol{h}_j, \|\vec{\boldsymbol{x}}_i - \vec{\boldsymbol{x}}_j\|^2, \boldsymbol{e}_{ij}), && (24)\\
\vec{\boldsymbol{x}}_i' &= \vec{\boldsymbol{x}}_i + \sum\nolimits_{j\in\mathcal{N}(i)}(\vec{\boldsymbol{x}}_i - \vec{\boldsymbol{x}}_j)\phi_x(\boldsymbol{m}_{ij}), && (25)\\
\vec{\boldsymbol{h}}_i' &= \phi_h(\boldsymbol{h}_i, \sum\nolimits_{j\in\mathcal{N}(i)}\boldsymbol{m}_{ij}), && (26)
\end{aligned}$$

where $\mathcal{N}(i)$ denotes the neighborhood of node $i$, and $\phi_x$ outputs a scalar. Therefore, for the invariant part, we have:

$$\begin{aligned}
f_i(\{\boldsymbol{h}_i, F_g(g\cdot\vec{\boldsymbol{x}}_i)\}) &= \phi_h(\boldsymbol{h}_i, \sum\nolimits_{j\in\mathcal{N}(i)}\boldsymbol{m}_{ij,F_g}) && (27)\\
&= \phi_h(\boldsymbol{h}_i, \sum\nolimits_{j\in\mathcal{N}(i)}\phi_m(\boldsymbol{h}_i, \boldsymbol{h}_j, \|F_g(g\cdot\vec{\boldsymbol{x}}_i) - F_g(g\cdot\vec{\boldsymbol{x}}_j)\|^2, \boldsymbol{e}_{ij})) && (28)\\
&= \phi_h(\boldsymbol{h}_i, \sum\nolimits_{j\in\mathcal{N}(i)}\phi_m(\boldsymbol{h}_i, \boldsymbol{h}_j, \|F(\vec{\boldsymbol{x}}_i) - F(\vec{\boldsymbol{x}}_j)\|, \boldsymbol{e}_{ij})) && (29)\\
&= \phi_h(\boldsymbol{h}_i, \sum\nolimits_{j\in\mathcal{N}(i)}\boldsymbol{m}_{ij,F}) && (30)\\
&= f_i(\{\boldsymbol{h}_i, F(\vec{\boldsymbol{x}}_i)\}) && (31)
\end{aligned}$$

For the equivariant features, recalling $F^{-1}(\vec{\boldsymbol{x}}) = \vec{\boldsymbol{L}}\vec{\boldsymbol{x}} + \mathbb{E}[\vec{\boldsymbol{X}}]$, we have:

$$
\begin{aligned}
& F_g^{-1}\vec{f}_i(\{\boldsymbol{h}_i, F_g(g\cdot\vec{\boldsymbol{x}}_i)\}) && (32)\\
&= F_g^{-1}(F_g(g\cdot\vec{\boldsymbol{x}}_i) + \sum\nolimits_{j\in\mathcal{N}(i)}(F_g(g\cdot\vec{\boldsymbol{x}}_i) - F_g(g\cdot\vec{\boldsymbol{x}}_j))\phi_x(\boldsymbol{m}_{ij,F_g})) && (33)\\
&= F_g^{-1}(F_g(g\cdot\vec{\boldsymbol{x}}_i) + \sum\nolimits_{j\in\mathcal{N}(i)}\vec{\boldsymbol{L}}_g^{-1}\boldsymbol{Q}(\vec{\boldsymbol{x}}_i - \vec{\boldsymbol{x}}_j)\phi_x(\boldsymbol{m}_{ij,F})) && (34)\\
&= F_g^{-1}(\vec{\boldsymbol{L}}_g^{-1}(\boldsymbol{Q}\vec{\boldsymbol{x}}_i + \vec{\boldsymbol{t}} - \mathbb{E}[g\cdot\vec{\boldsymbol{X}}]) + \sum\nolimits_{j\in\mathcal{N}(i)}\vec{\boldsymbol{L}}_g^{-1}\boldsymbol{Q}(\vec{\boldsymbol{x}}_i - \vec{\boldsymbol{x}}_j)\phi_x(\boldsymbol{m}_{ij,F})) && (35)\\
&= F_g^{-1}(\vec{\boldsymbol{L}}_g^{-1}(\boldsymbol{Q}\vec{\boldsymbol{x}}_i + \vec{\boldsymbol{t}} - g\cdot\mathbb{E}[\vec{\boldsymbol{X}}]) + \sum\nolimits_{j\in\mathcal{N}(i)}\vec{\boldsymbol{L}}_g^{-1}\boldsymbol{Q}(\vec{\boldsymbol{x}}_i - \vec{\boldsymbol{x}}_j)\phi_x(\boldsymbol{m}_{ij,F})) && (36)\\
&= F_g^{-1}(\vec{\boldsymbol{L}}_g^{-1}(\boldsymbol{Q}\vec{\boldsymbol{x}}_i - \boldsymbol{Q}\mathbb{E}[\vec{\boldsymbol{X}}]) + \sum\nolimits_{j\in\mathcal{N}(i)}\vec{\boldsymbol{L}}_g^{-1}\boldsymbol{Q}(\vec{\boldsymbol{x}}_i - \vec{\boldsymbol{x}}_j)\phi_x(\boldsymbol{m}_{ij,F})) && (37)\\
&= F_g^{-1}(\vec{\boldsymbol{L}}_g^{-1}\boldsymbol{Q}(\vec{\boldsymbol{x}}_i - \mathbb{E}[\vec{\boldsymbol{X}}]) + \sum\nolimits_{j\in\mathcal{N}(i)}\vec{\boldsymbol{L}}_g^{-1}\boldsymbol{Q}(\vec{\boldsymbol{x}}_i - \vec{\boldsymbol{x}}_j)\phi_x(\boldsymbol{m}_{ij,F})) && (38)\\
&= F_g^{-1}(\vec{\boldsymbol{L}}_g^{-1}\boldsymbol{Q}(\vec{\boldsymbol{x}}_i - \mathbb{E}[\vec{\boldsymbol{X}}] + \sum\nolimits_{j\in\mathcal{N}(i)}(\vec{\boldsymbol{x}}_i - \vec{\boldsymbol{x}}_j)\phi_x(\boldsymbol{m}_{ij,F}))) && (39)\\
&= \boldsymbol{Q}(\vec{\boldsymbol{x}}_i - \mathbb{E}[\vec{\boldsymbol{X}}] + \sum\nolimits_{j\in\mathcal{N}(i)}(\vec{\boldsymbol{x}}_i - \vec{\boldsymbol{x}}_j)\phi_x(\boldsymbol{m}_{ij,F})) + \mathbb{E}[g\cdot\vec{\boldsymbol{X}}] && (40)\\
&= \boldsymbol{Q}(\vec{\boldsymbol{x}}_i + \sum\nolimits_{j\in\mathcal{N}(i)}(\vec{\boldsymbol{x}}_i - \vec{\boldsymbol{x}}_j)\phi_x(\boldsymbol{m}_{ij,F})) - \boldsymbol{Q}\mathbb{E}[\vec{\boldsymbol{X}}] + g\cdot\mathbb{E}[\vec{\boldsymbol{X}}] && (41)\\
&= \boldsymbol{Q}(\vec{\boldsymbol{x}}_i + \sum\nolimits_{j\in\mathcal{N}(i)}(\vec{\boldsymbol{x}}_i - \vec{\boldsymbol{x}}_j)\phi_x(\boldsymbol{m}_{ij,F})) + \vec{\boldsymbol{t}} && (42)\\
&= g\cdot(\vec{\boldsymbol{x}}_i + \sum\nolimits_{j\in\mathcal{N}(i)}(\vec{\boldsymbol{x}}_i - \vec{\boldsymbol{x}}_j)\phi_x(\boldsymbol{m}_{ij,F})) && (43)
\end{aligned}
$$

By replacing $g$ with identical element $I$ in $E(3)$, since $F = F_I$, we can immediately derive:

$$F^{-1}\vec{f}_i((\{\boldsymbol{h}_i, F(\vec{\boldsymbol{x}}_i)\}) = (\vec{\boldsymbol{x}}_i + \sum\nolimits_{j\in\mathcal{N}(i)}(\vec{\boldsymbol{x}}_i - \vec{\boldsymbol{x}}_j)\phi_x(\boldsymbol{m}_{ij,F})), \quad (44)$$

$$g\cdot F^{-1}\vec{f}_i((\{\boldsymbol{h}_i, F(\vec{\boldsymbol{x}}_i)\}) = F_g^{-1}\vec{f}_i(\{\boldsymbol{h}_i, F_g(g\cdot\vec{\boldsymbol{x}}_i)\}), \quad (45)$$

which concludes Proposition 1.1. □

##### 1.3.3 Latent Diffusion Model

With the aforementioned preparations, the discrete residue types are encoded as continuous latent representations $\{\boldsymbol{z}_i\}$, and the all-atom geometry is also compressed and standardized into 3D vectors $\{\vec{\boldsymbol{z}}_i\} \sim \mathcal{N}(\boldsymbol{0}, \boldsymbol{I})$. Therefore, we are ready to implement a diffusion model on the standard latent space to generate $\boldsymbol{z}_i$ and $\vec{\boldsymbol{z}}_i$ conditioned on the binding site $\mathcal{G}_b$. The forward diffusion process gradually adds noise to the data from $t = 0$ to $t = T$, resulting in the prior distribution $\mathcal{N}(\boldsymbol{0}, \boldsymbol{I})$. The reverse diffusion process generates data distribution by iteratively denosing the distribution from $t = T$ to $t = 0$. We denote $\vec{\boldsymbol{u}}_i^t = [\boldsymbol{z}_i^t, \vec{\boldsymbol{z}}_i^t]$ and $\mathcal{G}_z^t = \{(\boldsymbol{z}_i^t, \vec{\boldsymbol{z}}_i^t)\}$ as the intermediate state for node $i$ and the entire peptide at time step $t$, respectively. For simplicity, we assume both $\mathcal{G}_z^t$ and the binding site $\mathcal{G}_b$ are already standardized via the transformation $F$ in Eq. 6. Then we have the forward process as:

$$q(\vec{\boldsymbol{u}}_i^t|\vec{\boldsymbol{u}}_i^{t-1}) = \mathcal{N}(\vec{\boldsymbol{u}}_i^t; \sqrt{1-\beta^t}\cdot\vec{\boldsymbol{u}}_i^{t-1}, \beta^t\boldsymbol{I}), \quad (46)$$

$$q(\vec{\boldsymbol{u}}_i^t|\vec{\boldsymbol{u}}_i^0) = \mathcal{N}(\vec{\boldsymbol{u}}_i^t; \sqrt{\bar{\alpha}^t}\cdot\vec{\boldsymbol{u}}_i^0, (1-\bar{\alpha}^t)\boldsymbol{I}), \quad (47)$$

where $\beta^t$ is the noise scale increasing with the timestep from 0 to 1 conforming to the cosine schedule[18], and $\bar{\alpha}^t = \prod_{s=1}^{s=t}(1-\beta^s)$. Then the state at timestep $t$ can be sampled as:

$$\vec{\boldsymbol{u}}_i^t = \sqrt{\bar{\alpha}^t}\vec{\boldsymbol{u}}_i^0 + (1-\bar{\alpha}^t)\boldsymbol{\epsilon}_i, \tag{48}$$

where $\boldsymbol{\epsilon}_i \sim \mathcal{N}(\mathbf{0}, \boldsymbol{I})$. Following[9], the reverse process can be defined with the reparameterization trick as:

$$p_\theta(\vec{\boldsymbol{u}}_i^{t-1}|\mathcal{G}_z^t, \mathcal{G}_b) = \mathcal{N}(\vec{\boldsymbol{u}}_i^{t-1}; \vec{\boldsymbol{\mu}}_\theta(\mathcal{G}_z^t, \mathcal{G}_b), \beta^t \boldsymbol{I}), \tag{49}$$

$$\vec{\boldsymbol{\mu}}_\theta(\mathcal{G}_z^t, \mathcal{G}_b) = \frac{1}{\sqrt{\alpha^t}}(\vec{\boldsymbol{u}}_i^t - \frac{\beta^t}{\sqrt{1-\bar{\alpha}^t}}\boldsymbol{\epsilon}_\theta(\mathcal{G}_z^t, \mathcal{G}_b, t)[i]), \tag{50}$$

where $\alpha^t = 1 - \beta^t$, and $\boldsymbol{\epsilon}_\theta$ is the denoising network also implemented with the adaptive multi-channel equivariant graph neural network (Supplementary Notes 1.3.5) to retain all-atom context of the binding site during generation and preserve the equivariance under affine transformations (Proposition 1.1). The training objective is illustrated in Supplementary Notes 1.4.2, and the training procedure is described in Supplementary Notes 1.5.2.

##### 1.3.4 Latent Interface Encoder

The latent interface encoder $\mathcal{E}_\mathcal{I}$ takes the latent point clouds at time step $t$ ($\mathcal{G}_z^t$) as input, and output a single numerical embedding of the interface $\boldsymbol{h}_\mathcal{G}^t$. To better processing the geometry of the latent point clouds, we parameterize the interface encoder with a geometric equivariant transformer (Algorithm 4). The training objective is to distinguishing similar interfaces from dissimilar ones (Supplementary Notes 1.4.3). In this way, the interface representation is equipped with the capabiliy of guiding the generation of peptides towards the directions that more resemble the reference interface (Supplementary Notes 1.6.2). Detailed algorithms for constructing the similar/dissimilar interface pairs and training the interface encoder are provided in Supplementary Notes 1.2.2 and 1.5.3, respectively.

##### 1.3.5 Algorithm for Adaptive Multi-Channel Equivariant Graph Neural Network

The adaptive multi-channel equivariant graph neural network (Algorithm 3) is adopted as the backbone model in both the all-atom variational autoencoder (Supplementary Notes 1.3.1) and the denoising network of the latent diffusion model (Supplementary Notes 1.3.3). Such choice originated from the requirements of capturing the all-atom structures where the number of atoms varies in different nodes with different amino acid types. While existing multi-channel graph neural networks (GNNs)[3,14] can only process nodes with fixed number of channels, the adaptive multi-channel equivariant GNN used here successfully tackles the variability of the channel size.

**Algorithm 3** Adaptive Multi-Channel Equivariant Graph Neural Network

**def** GeometricRelationExtractor($\vec{\boldsymbol{X}}_i, \vec{\boldsymbol{X}}_j$):

\# *Calculate Distance Matrix*

$\boldsymbol{D}_{ij}(p,q) \leftarrow ||\boldsymbol{X}_i[:,p] - \boldsymbol{X}_j[:,q]||_2$

\# *Extract Channel-wise Relations via Weighted Distances*

$\boldsymbol{R}_{ij} \leftarrow \boldsymbol{A}_i^\top(\boldsymbol{w}_i\boldsymbol{w}_j^\top \odot \boldsymbol{D}_{ij})\boldsymbol{A}_j$

**return** $\boldsymbol{R}_{ij}$

**def** GeometricMessageScaler($\vec{\boldsymbol{X}}, \boldsymbol{s}$):

\# *Channel-wise Convolution to Reshape C-Channel Scalers into c Channels*

$\boldsymbol{s}'[i] \leftarrow \frac{1}{C-c+1}\sum_{j=i-(C-c)/2}^{i+(C-c)/2} \boldsymbol{s}[j]$

\# *Multiply Reshaped Scalers on c-Channel Vectors*

$\boldsymbol{X}' \leftarrow \boldsymbol{X} \cdot \mathrm{diag}(\boldsymbol{s}')$

**return** $\boldsymbol{X}'$

**def** DynamicMultiChannelMessagePassingEncoder($\boldsymbol{H}, \vec{\mathbf{X}}$):

\# *Initialization*

$\boldsymbol{H}^{(0)}, \vec{\mathbf{X}}^{(0)} \leftarrow \boldsymbol{H}, \vec{\mathbf{X}}$

\# *Message Passing Loop*

**for** $l$ in $0, 1, \cdots, L-1$ **do**

$\boldsymbol{h}_i^{(l)}, \boldsymbol{h}_j^{(l)}, \vec{\boldsymbol{X}}_i^{(l)}, \vec{\boldsymbol{X}}_j^{(l)} \leftarrow \boldsymbol{H}^{(l)}[i], \boldsymbol{H}^{(l)}[j], \vec{\mathbf{X}}^{(l)}[i], \vec{\mathbf{X}}^{(l)}[j]$

$\boldsymbol{R}_{ij}^{(l)} \leftarrow$ GeometricRelationExtractor$(\vec{\boldsymbol{X}}_i^{(l)}, \vec{\boldsymbol{X}}_j^{(l)})$

$\boldsymbol{m}_{ij} \leftarrow \varphi_m\left(\boldsymbol{h}_i^{(l)}, \boldsymbol{h}_j^{(l)}, \frac{\boldsymbol{R}_{ij}^{(l)}}{||\boldsymbol{R}_{ij}^{(l)}||_F+\epsilon}\right)$

$\vec{\boldsymbol{X}}_{ij} \leftarrow$ GeometricMessageScaler$(\vec{\boldsymbol{X}}_i^{(l)} - \frac{1}{c_j}\sum_{k=1}^{c_j}\vec{\boldsymbol{X}}_j^{(l)}[:,k], \phi_x(\boldsymbol{m}_{ij}))$

$\boldsymbol{h}_i^{(l+1)} \leftarrow \varphi_h(\boldsymbol{h}_i^{(l)}, \sum_{j\in\mathcal{N}(i)}\boldsymbol{m}_{ij})$

$\vec{\boldsymbol{X}}_i^{(l+1)} \leftarrow \vec{\boldsymbol{X}}_i^{(l)} + \frac{1}{|\mathcal{N}(i)|}\sum_{j\in\mathcal{N}(i)}\vec{\boldsymbol{X}}_{ij}$

$\boldsymbol{H}^{(l+1)}, \vec{\mathbf{X}}^{(l+1)} \leftarrow \{\boldsymbol{h}_i^{(l+1)}\}_{i=1}^N, \{\vec{\boldsymbol{X}}_i^{(l+1)}\}_{i=1}^N$

**end for**

**return** $\boldsymbol{H}^{(L)}, \vec{\mathbf{X}}^{(L)}$

###### 1.3.6 Algorithm for Geometric Equivariant Transformer

The geometric equivariant transformer (Algorithm 4) is used in the latent interface encoder to better capture the overall geometry of the latent point clouds and compress them into single representations. The transformer architecture is known for its strong capacity in learning the precise representations of the data[12].

**Algorithm 4** Geometric Equivariant Transformer

**def** GraphEmbedding($\boldsymbol{H}, \vec{\boldsymbol{X}}$):
### *One-Layer Message Passing*
  $\boldsymbol{h}_j, \boldsymbol{h}_j, \vec{\boldsymbol{x}}_i, \vec{\boldsymbol{x}}_j \leftarrow \boldsymbol{H}[i], \boldsymbol{H}[j], \vec{\boldsymbol{X}}[i], \vec{\boldsymbol{X}}[j]$
  $\boldsymbol{m}_{ij} \leftarrow [\boldsymbol{h}_i, \boldsymbol{h}_j, \mathrm{RBF}(\|\vec{\boldsymbol{x}}_i - \vec{\boldsymbol{x}}_j\|_2)]$
  $\boldsymbol{h}_i \leftarrow \varphi_h(\boldsymbol{h}_i, \sum_{j \in \mathcal{N}(i)} \varphi_s(\boldsymbol{m}_{ij}) \cdot \boldsymbol{h}_j)$
  $\vec{\boldsymbol{v}}_i \leftarrow \sum_{j \in \mathcal{N}(i)} \varphi_v(\boldsymbol{m}_{ij}) \cdot (\vec{\boldsymbol{x}}_i - \vec{\boldsymbol{x}}_j)$
  $\boldsymbol{H}, \vec{\boldsymbol{V}} \leftarrow \{\boldsymbol{h}_i\}_{i=1}^N, \{\vec{\boldsymbol{v}}_i\}_{i=1}^N$
  **return** $\boldsymbol{H}, \vec{\boldsymbol{V}}$

**def** EquivariantSelfAttention($\boldsymbol{H}, \vec{\boldsymbol{V}}, \vec{\boldsymbol{X}}$):
### *Calculate Queries, Keys, and Values of the s-th Head*
  $\boldsymbol{Q}_s, \boldsymbol{K}_s, \boldsymbol{V}_s \leftarrow \boldsymbol{H}\boldsymbol{W}_s^Q, \boldsymbol{H}\boldsymbol{W}_s^K, [\boldsymbol{H}\boldsymbol{W}_s^{Vh}, \vec{\boldsymbol{V}}\boldsymbol{W}_s^{Vv}]$
### *Distance-Enhanced Self-Attention Matrix*
  $\boldsymbol{D} \leftarrow \{\vec{\boldsymbol{X}}[i] - \vec{\boldsymbol{X}}[j]\}_{i,j=1}^N$
  $\boldsymbol{R} \leftarrow \{\varphi_r(\mathrm{RBF}(\boldsymbol{D}_{ij}))\}_{j \in \mathcal{N}(i)}$
  $\boldsymbol{H}_s, \vec{\boldsymbol{V}}_s \leftarrow \mathrm{Softmax}\left(\frac{\mathbf{Q}_\mathbf{s}^\top \mathbf{K}_\mathbf{s}}{2\sqrt{h_s}} - \boldsymbol{D} + \boldsymbol{R}\right)\boldsymbol{V}_s$
### *Output Projection*
  $\boldsymbol{H}, \vec{\boldsymbol{V}} \leftarrow \sum_s \boldsymbol{H}_s \boldsymbol{W}_s^{Oh}, \sum_s \vec{\boldsymbol{V}}_s \boldsymbol{W}_s^{Ov}$
  **return** $\boldsymbol{H}, \vec{\boldsymbol{V}}$

**def** EquivariantFFN($\boldsymbol{H}, \vec{\boldsymbol{V}}$):
### *GVP-Style Feature Mixing*
  $\vec{\boldsymbol{V}}_1, \vec{\boldsymbol{V}}_2 \leftarrow \vec{\boldsymbol{V}}\boldsymbol{W}_1, \vec{\boldsymbol{V}}\boldsymbol{W}_2$
  $\boldsymbol{H}, \boldsymbol{U} \leftarrow \varphi_{\mathrm{FFN}}(\boldsymbol{H}, \|\vec{\boldsymbol{V}}_1\|_2)$
  $\vec{\boldsymbol{V}} \leftarrow \mathrm{LN}(\boldsymbol{U}) \odot \vec{\boldsymbol{V}}_2$
  **return** $\boldsymbol{H}, \vec{\boldsymbol{V}}$

**def** InterfaceEncoder($\boldsymbol{H}, \vec{\boldsymbol{X}}$):
### *Initialization by Graph Embedding Layer*
  $\boldsymbol{H}^{(0)}, \vec{\boldsymbol{V}}^{(0)} \leftarrow \mathrm{GraphEmbedding}(\boldsymbol{H}, \vec{\boldsymbol{X}})$
### *Message Passing Loop*
  **for** $l$ in $0, 1, \cdots, L-1$ **do**
### *Pre-Layer Normalization*
    $[\boldsymbol{H}^{(l)}, \vec{\boldsymbol{V}}^{(l)}] \leftarrow [\boldsymbol{H}^{(l)}, \vec{\boldsymbol{V}}^{(l)}] + \mathrm{EquivariantSelfAttention}(\mathrm{LayerNorm}(\boldsymbol{H}^{(l)}), \vec{\boldsymbol{V}}^{(l)}, \vec{\boldsymbol{X}})$
    $[\boldsymbol{H}^{(l+1)}, \vec{\boldsymbol{V}}^{(l+1)}] \leftarrow [\boldsymbol{H}^{(l)}, \vec{\boldsymbol{V}}^{(l)}] + \mathrm{EquivariantFFN}(\mathrm{LayerNorm}(\boldsymbol{H}^{(l)}), \vec{\boldsymbol{V}}^{(l)})$
  **end for**
### *Final Pooling*
  $\boldsymbol{h}_\mathcal{G} \leftarrow \varphi_\mathcal{G}(\frac{1}{\sqrt{N}}\mathbf{1}^\top \boldsymbol{H}^{(L)})$
  **return** $\boldsymbol{h}_\mathcal{G}$

##### 1.4 Training Objectives

###### 1.4.1 Losses for the All-Atom Variational AutoEncoder

As a Variational Autoencoder(VAE), the training objective consists of two key components, namely the reconstruction loss $\mathcal{L}_{recon}$ and the KL-divergence term $\mathcal{L}_{KL}$. For the reconstruction loss, the goal is to recover the encoded amino acid types and all-atom structures of each node $i$, which is detailed as

$$\mathcal{L}_{recon}(i) = \text{CrossEntropy}(p(x_i), p(x_i')) + \text{MSE}(\vec{\boldsymbol{X}}_i, \vec{\boldsymbol{X}}_i') + \mathcal{L}_{aux}(i), \tag{51}$$

where the auxilary loss $\mathcal{L}_{aux}$ is applied for more precise all-atom reconstruction with supervision on $\mathtt{C}_\alpha$ coordinates, bond lengths, and side-chain dihedral angles. First, since the $\mathtt{C}_\alpha$ is critical in deciding the global geometry of the peptide, we exert additional loss on its coordinates to distinguish it from other atoms:

$$\mathcal{L}_{CA}(i) = \text{MSE}(\vec{\boldsymbol{r}}_i, \vec{\boldsymbol{r}}_i'), \tag{52}$$

where $\vec{\boldsymbol{r}}_i'$ and $\vec{\boldsymbol{r}}_i$ are the reconstructed and the ground truth coordinates of $\mathtt{C}_\alpha$ in node $i$. Next, we implement L1 loss on the bond lengths:

$$\mathcal{L}_{bond}(i) = \sum\nolimits_{b \in \mathcal{B}(i)} |b - b'| / |\mathcal{B}(i)|, \tag{53}$$

where $\mathcal{B}(i)$ includes all chemical bonds in node $i$, and $b'$ denotes the reconstructed bond length. For simplicity, he bonds between residues are included into the bonds of the former residue. Finally, we supervise on the $\chi_1$ to $\chi_4$ side-chain dihedral angles:

$$\mathcal{L}_{angle}(i) = \sum\nolimits_{\chi \in \mathcal{A}(i)} |\chi - \chi'| / |\mathcal{A}(i)|, \tag{54}$$

where $\mathcal{A}(i)$ includes all side-chain dihedral angles in node $i$, $\chi'$ and $\chi$ denotes the reconstructed and the ground truth angles, respectively. The overall auxilary loss for node $i$ is then given by:

$$\mathcal{L}_{aux}(i) = \lambda_{CA}\mathcal{L}_{CA}(i) + \lambda_{bond}\mathcal{L}_{bond}(i) + \lambda_{angle}\mathcal{L}_{angle}(i), \tag{55}$$

where we set $\lambda_{CA} = 1.0, \lambda_{bond} = 1.0, \lambda_{angle} = 0.5$ in our experiments. We find that it is necessary to set $\lambda_{angle}$ with a relatively small value to make the training process stable. Finally, the KL-divergence prevents the scale of $\boldsymbol{z}_i$ from exploding and constrains $\vec{\boldsymbol{z}}_i$ around $\mathtt{C}_\alpha$ to retain necessary geometric information:

$$\begin{aligned} \mathcal{L}_{KL}(i) = \quad & \lambda_1 \cdot D_{\text{KL}}(\mathcal{N}(\boldsymbol{0}, \boldsymbol{I}) \| \mathcal{N}(\boldsymbol{\mu}_i, \text{diag}(\boldsymbol{\sigma}_i))) \\ & + \lambda_2 \cdot D_{\text{KL}}(\mathcal{N}(\vec{\boldsymbol{r}}_i, \boldsymbol{I}) \| \mathcal{N}(\vec{\boldsymbol{\mu}}_i, \text{diag}(\vec{\boldsymbol{\sigma}}_i))), \end{aligned} \tag{56}$$

where $\lambda_1$ and $\lambda_2$ reweight the contraints on the sequence and the structure, respectively. And the overall training objective is:

$$\mathcal{L}_{AE} = \mathbb{E}_{\mathcal{G}_p}\Big[\sum\nolimits_{i \in \mathcal{G}_p} (\mathcal{L}_{recon}(i) + \mathcal{L}_{KL}(i)) / |\mathcal{G}_p|\Big]. \tag{57}$$

###### 1.4.2 Losses for the Latent Diffusion Model

The training objective of the latent diffusion model follows the standard DDPM[9] paradigm. The objective at time step $t$ is calculated as MSE between the predicted noise and the added noise in Eq. 48, and the overall training objective $\mathcal{L}_{LDM}$ is the expectation with respect to $t$:

$$\mathcal{L}_{LDM} = \mathbb{E}_{t \sim \mathbf{U}(1,T), \boldsymbol{\epsilon}_i \sim \mathcal{N}(0,1), \mathcal{G}_z^t, \mathcal{G}_b}\Big[\sum\nolimits_i \|\boldsymbol{\epsilon}_i - \boldsymbol{\epsilon}_\theta(\mathcal{G}_z^t, \mathcal{G}_b, t)[i]\|_2^2 / |\mathcal{G}_z^t|\Big], \tag{58}$$

where $\mathbf{U}(1,T)$ denotes uniform distribution of integers between 1 to $T$.

##### 1.4.3 Losses for the Latent Interface Encoder

Given the downsampled pairwise data, the target of the latent interface encoder is to minimize the L2-distance between the reference and the positive sample, and ensure the negative distance larger than a pre-defined threshold $\delta$. Specifically, we have

$$\mathcal{L}_{LDM} = \mathbb{E}_{t\sim \mathbf{U}(1,T),\mathcal{G}_z^t,\mathcal{G}_{z,+}^t,\mathcal{G}_{z,-}^t}\Big[\|\mathcal{E}_{\mathcal{I}}(\mathcal{G}_z^t) - \mathcal{E}_{\mathcal{I}}(\mathcal{G}_{z,+}^t)\|_2 + \max(\delta - \|\mathcal{E}_{\mathcal{I}}(\mathcal{G}_z^t) - \mathcal{E}_{\mathcal{I}}(\mathcal{G}_{z,-}^t)\|_2, 0)\Big]. \quad (59)$$

#### 1.5 Training Regimen

##### 1.5.1 Algorithm for Training All-Atom Variational AutoEncoder

The all-atom variational autoencoder was trained on the dataset merging the 70K peptide-like protein fragments and the 4k protein-peptide complexes (Methods). To encourage $\mathcal{E}_\phi$ to learn contextual representations of residues, we corrupt 25% of the residues in $\mathcal{G}_p$ with a [MASK] type to obtain $\tilde{\mathcal{G}}_p$ as the input. The overall procedure follows the Algorithm 5 depicted below.

**Algorithm 5** Training Algorithm of All-Atom Variational AutoEncoder

**def** TrainAutoEncoder($\mathcal{S}$):
  Initialize $\mathcal{E}_\phi$, $\mathcal{D}_\xi$
  **while** $\phi, \xi$ have not converged **do**
### *Sample from the dataset. In implementation, a batch of samples are drawn.*
    Sample $(\mathcal{G}_p, \mathcal{G}_b) \sim \mathcal{S}$
### *Mask 25% Residues*
    $\tilde{\mathcal{G}}_p \leftarrow \text{mask}(\mathcal{G}_p)$
### *Encoding*
    $\{(\boldsymbol{\mu}_i, \boldsymbol{\sigma}_i, \vec{\boldsymbol{\mu}}_i, \vec{\boldsymbol{\sigma}}_i)\} \leftarrow \mathcal{E}_\phi(\tilde{\mathcal{G}}_p, \mathcal{G}_b)$
### *Reparameterization*
    $\{(\boldsymbol{\epsilon}_i, \vec{\boldsymbol{\epsilon}}_i)\} \sim \mathcal{N}(\mathbf{0}, \boldsymbol{I})$
    $\mathcal{G}_z \leftarrow \{(\boldsymbol{\mu}_i + \boldsymbol{\epsilon}_i \odot \boldsymbol{\sigma}_i, \vec{\boldsymbol{\mu}}_i + \vec{\boldsymbol{\epsilon}}_i \odot \vec{\boldsymbol{\sigma}}_i)\}$
### *Decoding*
    $\mathcal{G}_p' \leftarrow \mathcal{D}_\xi(\mathcal{G}_z, \mathcal{G}_b)$
    $\mathcal{L}_{AE} = \sum_{i\in\mathcal{G}_p}(\mathcal{L}_{recon}(i) + \mathcal{L}_{KL}(i))/|\mathcal{G}_p|$
    $\phi, \xi \leftarrow \text{optimizer}(\mathcal{L}_{AE}; \phi, \xi)$
  **end while**
  **return** $\mathcal{E}_\phi$, $\mathcal{D}_\xi$

##### 1.5.2 Algorithm for Training Latent Diffusion

The training of the latent diffusion model were splitted into two stages: a pretraining stage on the 70K peptide-like protein fragments followed by a finetuning stage on the 4k protein-peptide complexes (Methods). Each stage of training follows Algorithm 6 described below.

**Algorithm 6** Training Algorithm of Latent Diffusion Model

**def** TrainLatentDiffusion($\mathcal{E}_\phi, \mathcal{D}_\xi, \mathcal{S}$):
  Initialize $\boldsymbol{\epsilon}_\theta$
  **while** $\theta$ have not converged **do**
    Sample $(\mathcal{G}_p, \mathcal{G}_b) \sim \mathcal{S}$
### *Encoding*
    $\{(\boldsymbol{\mu}_i, \boldsymbol{\sigma}_i, \vec{\boldsymbol{\mu}}_i, \vec{\boldsymbol{\sigma}}_i)\} \leftarrow \mathcal{E}_\phi(\mathcal{G}_p, \mathcal{G}_b)$
    $\mathcal{G}_z^0 \leftarrow \{(\boldsymbol{\mu}_i, \vec{\boldsymbol{\mu}}_i)\}$
### *Affine Transformation to Standard Geometry*
    $F \leftarrow \text{affine}(\mathcal{G}_b)$
    $\mathcal{G}_z^0, \mathcal{G}_b \leftarrow F(\mathcal{G}_z^0), F(\mathcal{G}_b)$
    $\{(\boldsymbol{z}_i^0, \vec{\boldsymbol{z}}_i^0)\}_{i=1}^N \leftarrow \mathcal{G}_z^0$
    $t \sim \mathbf{U}(1, T)$, $\{(\boldsymbol{\epsilon}_i, \vec{\boldsymbol{\epsilon}}_i)\}_{i=1}^N \sim \mathcal{N}(\mathbf{0}, \boldsymbol{I})$
    $\mathcal{G}_z^t \leftarrow \{\sqrt{\bar{\alpha}^t}[\boldsymbol{z}_i^0, \vec{\boldsymbol{z}}_i^0] + (1 - \bar{\alpha}^t)[\boldsymbol{\epsilon}_i, \vec{\boldsymbol{\epsilon}}_i]\}_{i=1}^N$
    $\mathcal{L}_{LDM}^t \leftarrow \sum_i \|[\boldsymbol{\epsilon}_i, \vec{\boldsymbol{\epsilon}}_i] - \boldsymbol{\epsilon}_\theta(\mathcal{G}_z^t, \mathcal{G}_b, t)[i]\|_2^2 / |\mathcal{G}_z^t|$
    $\theta \leftarrow \text{optimizer}(\mathcal{L}_{LDM}^t; \theta)$
  **end while**
  **return** $\boldsymbol{\epsilon}_\theta$

##### 1.5.3 Algorithm for Training Latent Interface Encoder

The latent interface encoder was also trained on the dataset merging the 70K peptide-like protein fragments and the 4k protein-peptide complexes (Methods), with the procedure illustrated below (Algorithm 7). The strategy for constructing positive (similar) and negative (dissimilar) interface pairs is demonstrated in Supplementary Notes 1.2.2.

**Algorithm 7** Training Algorithm of Latent Interface Encoder

**def** TimeDependentGraph($\mathcal{G}_z^0, t$):
### *Add Noise to Initial Graph towards Timestep t*
  $\{(\boldsymbol{z}_i^0, \vec{\boldsymbol{z}}_i^0)\}_{i=1}^N \leftarrow \mathcal{G}_z^0$
  $\{(\boldsymbol{\epsilon}_i, \vec{\boldsymbol{\epsilon}}_i)\}_{i=1}^N \sim \mathcal{N}(\boldsymbol{0}, \boldsymbol{I})$
  $\mathcal{G}_z^t \leftarrow \{\sqrt{\bar{\alpha}^t}[\boldsymbol{z}_i^0, \vec{\boldsymbol{z}}_i^0] + (1 - \bar{\alpha}^t)[\boldsymbol{\epsilon}_i, \vec{\boldsymbol{\epsilon}}_i]\}_{i=1}^N$
  **return** $\mathcal{G}_z^t$

**def** TrainInterface($\mathcal{S}, \mathcal{E}_\phi$):
  Initialize $\mathcal{E}_\mathcal{I}$
  **while** $\mathcal{I}$ have not converged **do**
    Sample $(\mathcal{G}_p, \mathcal{G}_b) \sim \mathcal{S}$
### *Encoding*
    $\{(\boldsymbol{\mu}_i, \boldsymbol{\sigma}_i, \vec{\boldsymbol{\mu}}_i, \vec{\boldsymbol{\sigma}}_i)\} \leftarrow \mathcal{E}_\phi(\mathcal{G}_p, \mathcal{G}_b)$
    $\mathcal{G}_z^0 \leftarrow \{(\boldsymbol{\mu}_i, \vec{\boldsymbol{\mu}}_i)\}$
### *Affine Transformation to Standard Geometry*
    $F \leftarrow \text{affine}(\mathcal{G}_b)$
    $\mathcal{G}_z^0 \leftarrow F(\mathcal{G}_z^0)$
### *Sample Pairwise Subgraphs via Down-Sampling*
    $\mathcal{G}_{z,\text{ref}}^0, \mathcal{G}_{z,+}^0, \mathcal{G}_{z,-}^0 \leftarrow \text{PairSampling}(\mathcal{G}_z^0)$
    $t \sim \mathbf{U}(1, T)$
    $\mathcal{G}_{z,\text{ref}}^t \leftarrow \text{TimeDependentGraph}(\mathcal{G}_{z,\text{ref}}^0, t)$
    $\mathcal{G}_{z,+}^t \leftarrow \text{TimeDependentGraph}(\mathcal{G}_{z,+}^0, t)$
    $\mathcal{G}_{z,-}^t \leftarrow \text{TimeDependentGraph}(\mathcal{G}_{z,-}^0, t)$
    $\boldsymbol{h}_{\mathcal{G},\text{ref}}^t, \boldsymbol{h}_{\mathcal{G},+}^t, \boldsymbol{h}_{\mathcal{G},-}^t \leftarrow \mathcal{E}_\mathcal{I}(\mathcal{G}_{z,\text{ref}}^t), \mathcal{E}_\mathcal{I}(\mathcal{G}_{z,+}^t), \mathcal{E}_\mathcal{I}(\mathcal{G}_{z,-}^t)$
    $\mathcal{L}_{IF}^t \leftarrow \|\boldsymbol{h}_{\mathcal{G},\text{ref}}^t - \boldsymbol{h}_{\mathcal{G},+}^t\|_2 + \max(\delta - \|\boldsymbol{h}_{\mathcal{G},\text{ref}}^t - \boldsymbol{h}_{\mathcal{G},-}^t\|_2, 0)$
    $\mathcal{I} \leftarrow \text{optimizer}(\mathcal{L}_{IF}^t; \mathcal{I})$
  **end while**
  **return** $\mathcal{E}_\mathcal{I}$

#### 1.6 Inference Regimen

##### 1.6.1 Algorithm for Inference

**Algorithm 8** Sampling Algorithm of PepMimic

**def** SampleFromPepMimic($\mathcal{D}_\xi, \boldsymbol{\epsilon}_\theta, \mathcal{G}_b$):

\# *Affine Transformation to Standard Geometry*

$F \leftarrow \text{affine}(\mathcal{G}_b)$

$\mathcal{G}_b \leftarrow F(\mathcal{G}_b)$

$\{(\boldsymbol{z}_i^T, \vec{\boldsymbol{z}}_i^T)\} \sim \mathcal{N}(\mathbf{0}, \boldsymbol{I})$

\# *Latent Denoising Loop*

**for** $t$ in $T, T-1, \cdots, 1$ **do**

$\quad \mathcal{G}_z^t \leftarrow \{(\boldsymbol{z}_i^t, \vec{\boldsymbol{z}}_i^t)\}$

$\quad \vec{\boldsymbol{u}}_i^t \leftarrow \boldsymbol{z}_i^t, \vec{\boldsymbol{z}}_i^t$

$\quad \boldsymbol{\varepsilon}_i = [\boldsymbol{\epsilon}_i, \vec{\boldsymbol{\epsilon}}_i] \sim \mathcal{N}(\mathbf{0}, \boldsymbol{I})$

$\quad \vec{\boldsymbol{u}}_i^{t-1} \leftarrow \frac{1}{\sqrt{\alpha^t}}(\vec{\boldsymbol{u}}_i^t - \frac{\beta^t}{\sqrt{1-\bar{\alpha}^t}}\boldsymbol{\epsilon}_\theta(\mathcal{G}_z^t, \mathcal{G}_b, t)[i]) + \beta_t \boldsymbol{\varepsilon}_i$

$\quad [\boldsymbol{z}_i^{t-1}, \vec{\boldsymbol{z}}_i^{t-1}] \leftarrow \vec{\boldsymbol{u}}_i^{t-1}$

**end for**

$\mathcal{G}_z \leftarrow \{(\boldsymbol{z}_i^0, \vec{\boldsymbol{z}}_i^0)\}$

\# *Back to Data Geometry*

$\mathcal{G}_z, \mathcal{G}_b \leftarrow F^{-1}(\mathcal{G}_z), F^{-1}(\mathcal{G}_b)$

\# *Decoding*

$\mathcal{G}_p \leftarrow \mathcal{D}_\xi(\mathcal{G}_z, \mathcal{G}_b)$

**return** $\mathcal{G}_p$

###### 1.6.2 Algorithm for Guided Diffusion

**Algorithm 9** Guided Sampling Algorithm of PepMimic

**def** GuidedSampleFromPepMimic($\mathcal{E}_\phi, \mathcal{E}_\mathcal{I}, \mathcal{D}_\xi, \boldsymbol{\epsilon}_\theta, \mathcal{G}_b, \mathcal{G}_r$):
\# *Encode Reference to Latent Space*
  $\{(\boldsymbol{\mu}_i, \boldsymbol{\sigma}_i, \vec{\boldsymbol{\mu}}_i, \vec{\boldsymbol{\sigma}}_i)\} \leftarrow \mathcal{E}_\phi(\mathcal{G}_r, \mathcal{G}_b)$
  $\mathcal{G}_r \leftarrow \{(\boldsymbol{\mu}_i, \vec{\boldsymbol{\mu}}_i)\}$
\# *Affine Transformation to Standard Geometry*
  $F \leftarrow \text{affine}(\mathcal{G}_b)$
  $\mathcal{G}_b \leftarrow F(\mathcal{G}_b)$
  $\{(\boldsymbol{z}_i^T, \vec{\boldsymbol{z}}_i^T)\} \sim \mathcal{N}(\boldsymbol{0}, \boldsymbol{I})$
\# *Latent Denoising Loop*
  **for** $t$ in $T, T-1, \cdots, 1$ **do**
    $\mathcal{G}_z^t \leftarrow \{(\boldsymbol{z}_i^t, \vec{\boldsymbol{z}}_i^t)\}$
\#   *Calculate Distance between Reference and Generated Sample*
    $d(\mathcal{G}_r, \mathcal{G}_z^t) \leftarrow \|\mathcal{E}_\mathcal{I}(\mathcal{G}_r) - \mathcal{E}_\mathcal{I}(\mathcal{G}_z^t)\|_2$
    $\vec{\boldsymbol{u}}_i^t \leftarrow \boldsymbol{z}_i^t, \vec{\boldsymbol{z}}_i^t$
    $\boldsymbol{\varepsilon}_i = [\boldsymbol{\epsilon}_i, \vec{\boldsymbol{\epsilon}}_i] \sim \mathcal{N}(\boldsymbol{0}, \boldsymbol{I})$
\#   *Rectify Denoising Terms via Gradients*
    $\vec{\boldsymbol{u}}_i^{t-1} \leftarrow \frac{1}{\sqrt{\alpha^t}}\left(\vec{\boldsymbol{u}}_i^t - \frac{\beta^t}{\sqrt{1-\bar{\alpha}^t}}(\boldsymbol{\epsilon}_\theta(\mathcal{G}_z^t, \mathcal{G}_b, t)[i] - \lambda\sqrt{1-\bar{\alpha}^t}\nabla_{\vec{\boldsymbol{u}}_i^t} d(\mathcal{G}_r, \mathcal{G}_z^t))\right) + \beta_t \boldsymbol{\varepsilon}_i$
    $[\boldsymbol{z}_i^{t-1}, \vec{\boldsymbol{z}}_i^{t-1}] \leftarrow \vec{\boldsymbol{u}}_i^{t-1}$
  **end for**
  $\mathcal{G}_z \leftarrow \{(\boldsymbol{z}_i^0, \vec{\boldsymbol{z}}_i^0)\}$
\# *Back to Data Geometry*
  $\mathcal{G}_z, \mathcal{G}_b \leftarrow F^{-1}(\mathcal{G}_z), F^{-1}(\mathcal{G}_b)$
\# *Decoding*
  $\mathcal{G}_p \leftarrow \mathcal{D}_\xi(\mathcal{G}_z, \mathcal{G}_b)$
  **return** $\mathcal{G}_p$

#### 2 Baseline Implementations

**HSRN**[10] We used the officially open-sourced codes (https://github.com/wengong-jin/abdockgen) for model construction, with hidden size = 128, and number of layers = 3. The model was trained with the same data and splits as our PepMimic, using AdamW[17] optimizer with learning rate = 1.0e-4, epoch = 50, and batch size = 8. The learning rate was decayed by 40% after five epochs without improvement on the validation set.

**dyMEAN**[15] We also used its official public codes (https://github.com/THUNLP-MT/dyMEAN) to construct the model, with hidden size = 128, number of layers = 3, and number of iterations = 3. Unlike antibodies, peptides were not constrainted by frameworks, thus we adjusted the algorithm to directly genenerate peptides at the epitopes, discarding the "shadow paratope"[15] mechanism. The model was trained with the same data and splits as our PepMimic, using AdamW[17] optimizer with learning rate = 1.0e-4, epoch = 100, and batch size = 32. The learning rate was decayed by 40% after five epochs without improvement on the validation set.

**RFDiffusion**[21] As it did not open-sourced the codes for training, we directly ran the official codes for inference (https://github.com/RosettaCommons/RFdiffusion) with the provided checkpoints for binder design. Following the official instruction (https://github.com/RosettaCommons/RFdiffusion?tab=readme-ov-file#binder-design), given a protein-peptide complex, we selected the longest continuous contig within 10Å $C_\beta$ distance to the peptide as the epitope, and randomly sampled 20% of the residues as hotspots for each generation. As instructed, with the backbone designed by RFDiffusion, we further used ProteinMPNN[2] for inverse folding to derive the sequence (https://github.com/nrbennet/dl_binder_design).

#### 3 Normalized Rosetta Interface Energy

Since interface energies between different binding sites are not directly comparable, we normalized the Rosetta interface energy[1] based on the energy distribution of candidates derived from each reference binding complex to evaluate its ability to distinguish successful designs from failed ones (Fig. 4g). Specifically, the normalization includes subtracting the average energy score of the candidates mimicking the same binding complex. This normalized Rosetta energy demonstrated certain capability of differentiating positive binders from negative ones (Fig. 4g).

#### 4 Implementation Details

**Generative Model** We trained PepMimic on a GPU with 24G memory with AdamW[17] optimizer. The autoencoder was trained on both the protein-peptide complexes and the augmentation fragments for 100 epochs with dynamic batches, ensuring that the total number of edges (proportional to the square of the number of nodes) remains below 60,000. The initial learning rate was $10^{-4}$, with a decay of 20% if the validation loss did not decrease for five consecutive epochs.

The diffusion model in the latent space was first pretrained on the augmented fragments for 100 epochs with a learning rate of $10^{-4}$. It was then fine-tuned on protein-peptide complexes for another 100 epochs, with an initial learning rate of $5 \times 10^{-5}$. During both pretraining and fine-tuning, the learning rate was decayed by 40% if the validation loss did not improve for three consecutive epochs, with validation conducted every 10 epochs. The batching strategy was consistent with that used for the autoencoder training.

The latent interface encoder was trained on both protein-peptide complexes and augmented fragments for 100 epochs with a learning rate of $10^{-4}$. The learning rate was decayed by 20% if the validation loss did not decrease for five consecutive epochs. The batching algorithm was similar to that used for training the autoencoder, except that the threshold for the total number of edges was increased to 480,000. Additional hyperparameters of PepMimic used in our experiments are provided in Supplementary Table 7.

**Ranking Strategy** For calculating rosetta energy[1], we utilized pyRosetta 4.0 with python 3.9, version 2023.49+release.9891f2c. Default REF2015 force fields were adopted for energy calculations. FoldX[19] was accessed through the official suite, which was downloaded from the FoldX website under an academic license (https://foldxsuite.crg.eu/products?page=1). For Alphafold Multimer[5,11], we used the codebase from the official Alphafold V2.3 repository (https://github.com/google-deepmind/alphafold), along with the parameters released on December 6th, 2022 (https://storage.googleapis.com/alphafold/alphafold_params_2022-12-06.tar). The genetic databases used were BFD (only version available), MGnify (v2022_05), UniRef90 (v2024_01), UniCluster30 (v2021_03), and UniProt (v2024_01). Template databases included PDB (downloaded on 2024-04-23) and PDB70 (2020-04-01). Multiple sequence alignments and template searches were conducted only for the target proteins, excluding the peptides. The number of cycles was set to the default value (i.e. 5).

#### Supplementary Figures

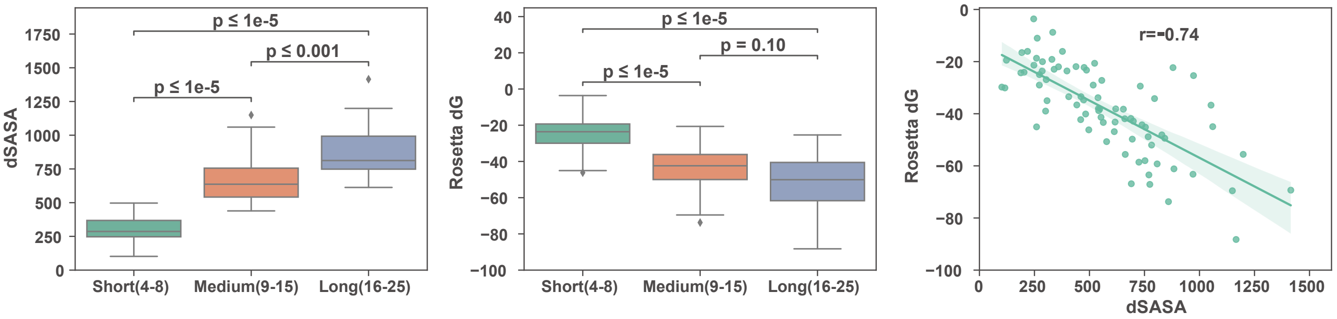

**Supplementary Figure 1. Solvent-accessible surface area (SASA) and Rosetta interface energy of the protein-peptide complexes in the test set split by lengths of the peptides.** **(a)** $\Delta$SASA ($\mathring{A}^2$) between unbounded state and bounded state. Longer peptides commonly form larger interfaces with higher $\Delta$SASA. **(b)** Rosetta interface energy ($\Delta G$) of the complexes. Longer peptides commonly achieve lower interface energy. **(c)** Pearson correlation between $\Delta$SASA and Rosetta $\Delta G$. The correlation is strong with a coefficient of $-0.74$ and a $p$-value $< 0.001$.

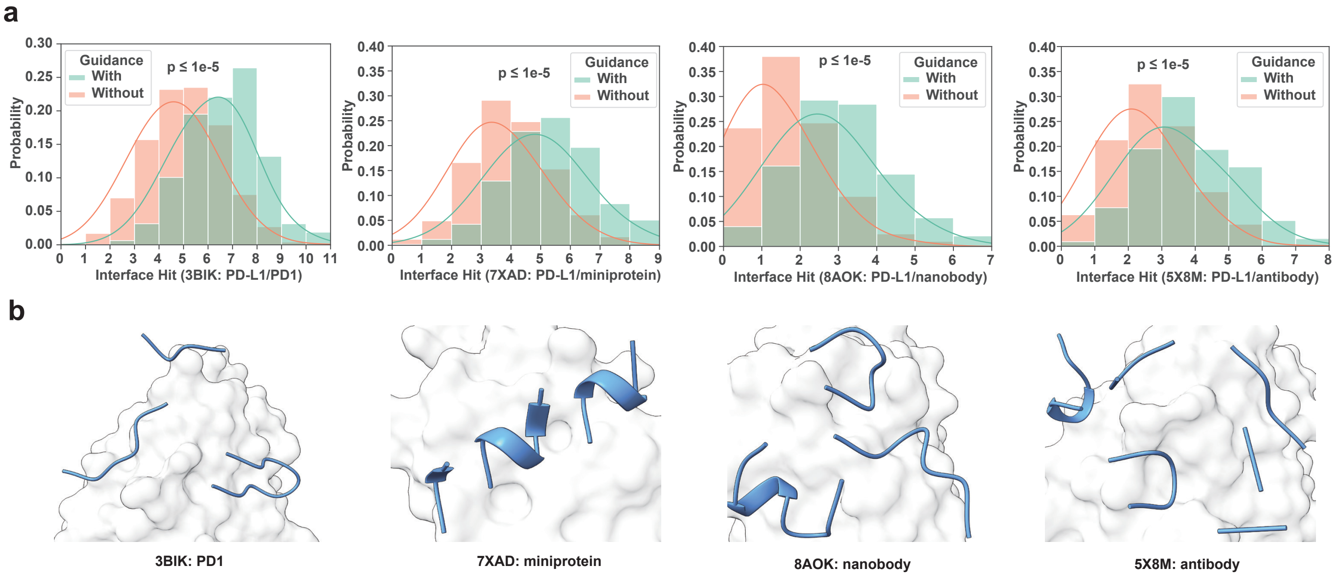

**Supplementary Figure 2.** **(a)** Distribution of interface hit for generated peptides with and without latent guidance. Results are categorized based on the type of the reference interface on PD-L1 formed by PD1, miniprotein, nanobody, and antibody. **(b)** Visualization of interfaces on PD-L1 with four different binding partners: PD1 (PDB ID 3BIK), AI-designed miniprotein (PDB ID 7XAD), nanobody (PDB ID 8AOK), and antibody durvalumab (PDB ID 5X8M).

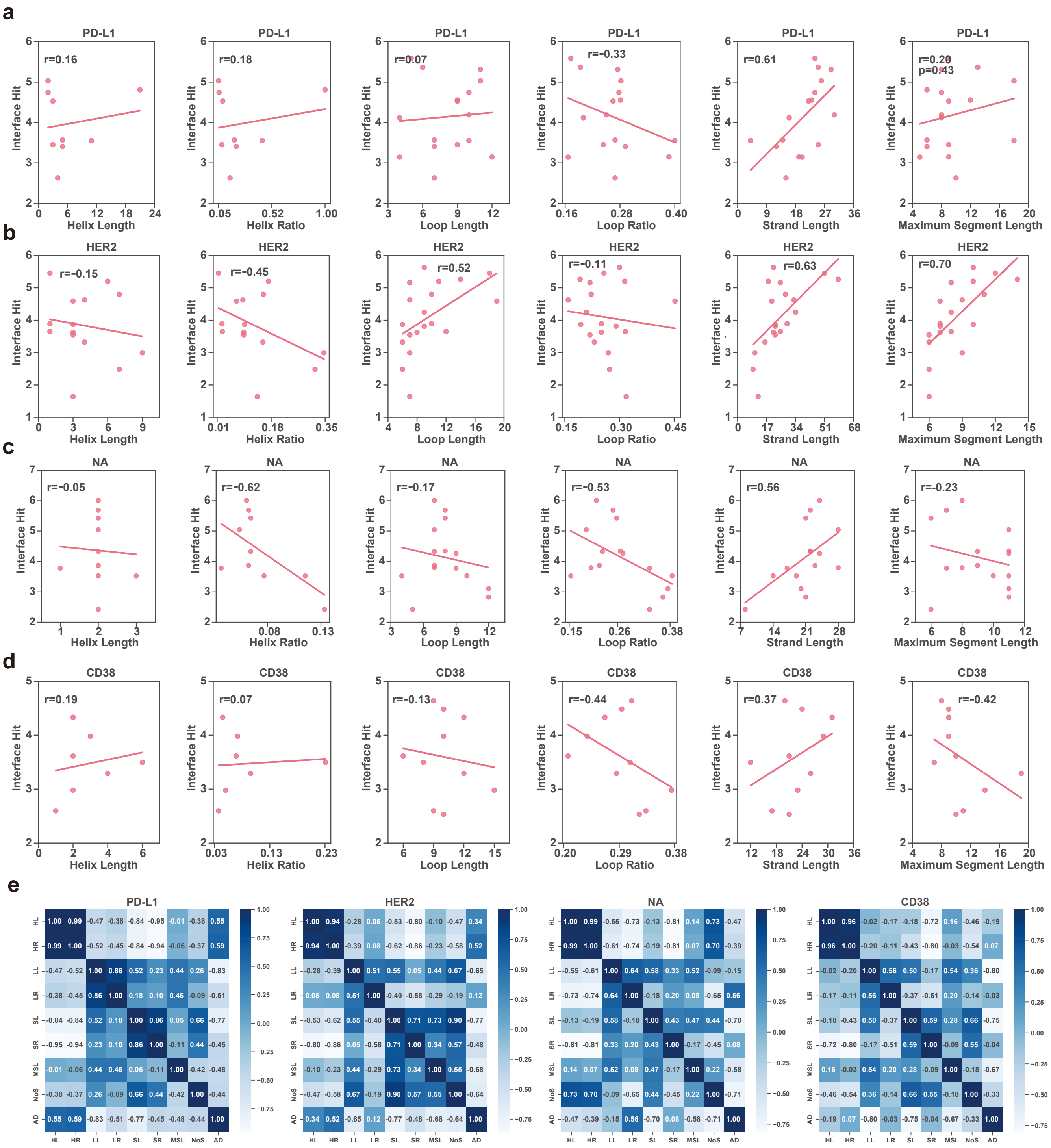

**Supplementary Figure 3.** **(a-d)** Pearson correlation between interface characteristics and interface hits of generated peptide mimicries, with each row of the results from PD-L1, HER2, viral neuraminidase (NA), and CD38, respectively. **(e)** Mutual Pearson correlations of the following characteristics on the interfaces: helix length (HL), helix ratio (HL), loop length (LL), loop ratio (LR), strand length (SL), strand ratio (SR), maximum segment length (MSL), number of segments (NoS), and average distance (AD) between interface residues.

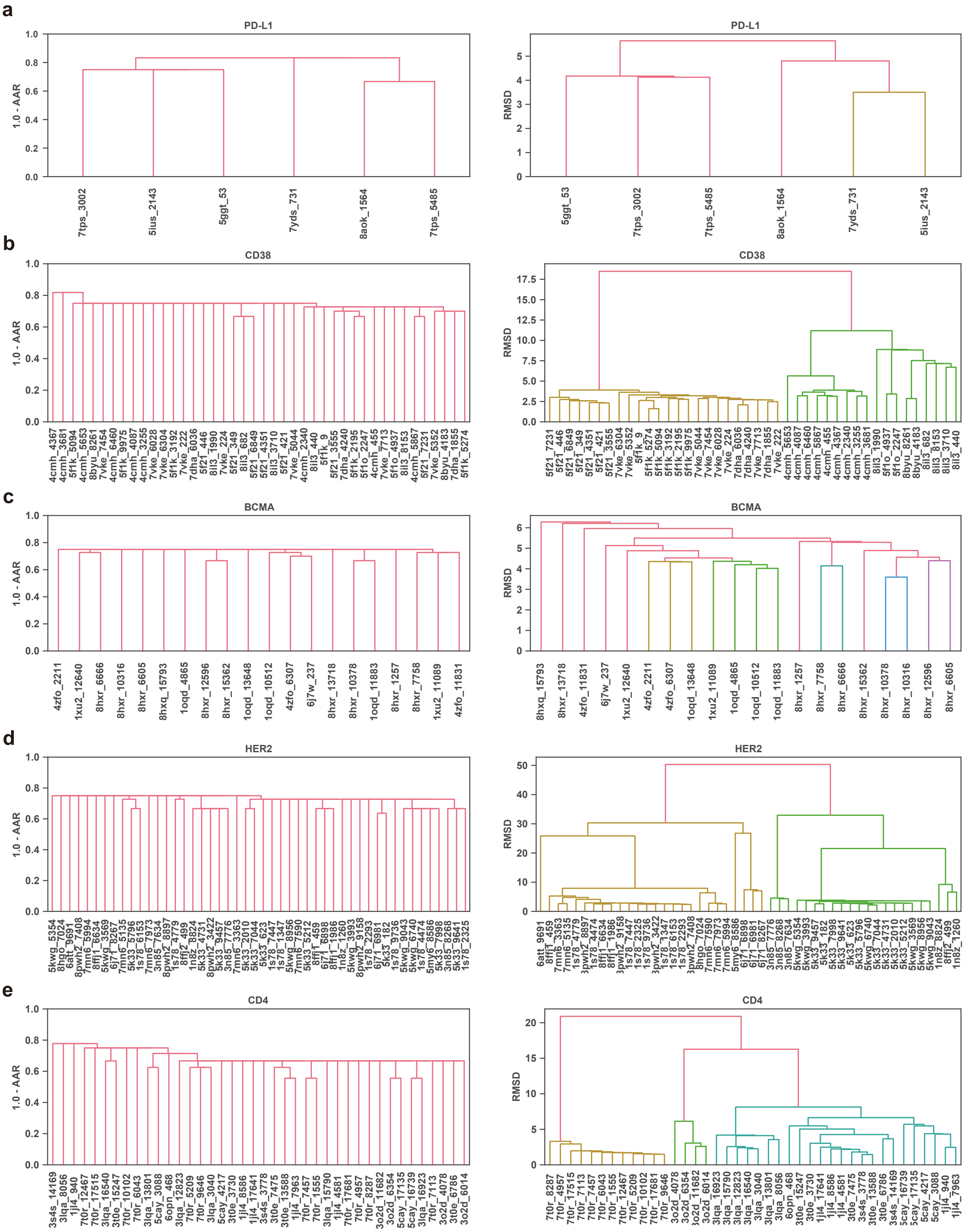

**Supplementary Figure 4.** **(a-e)** Sequence and structure hierarchical clusters of the experimentally successful designs ($K_D < 100$ nM). The distance metric for sequence is 1.0− Amino Acid Recovery (AAR), and the distance metric for structure is Root Mean Square Deviation (RMSD) of the peptide with coordinate system aligned by the target protein.

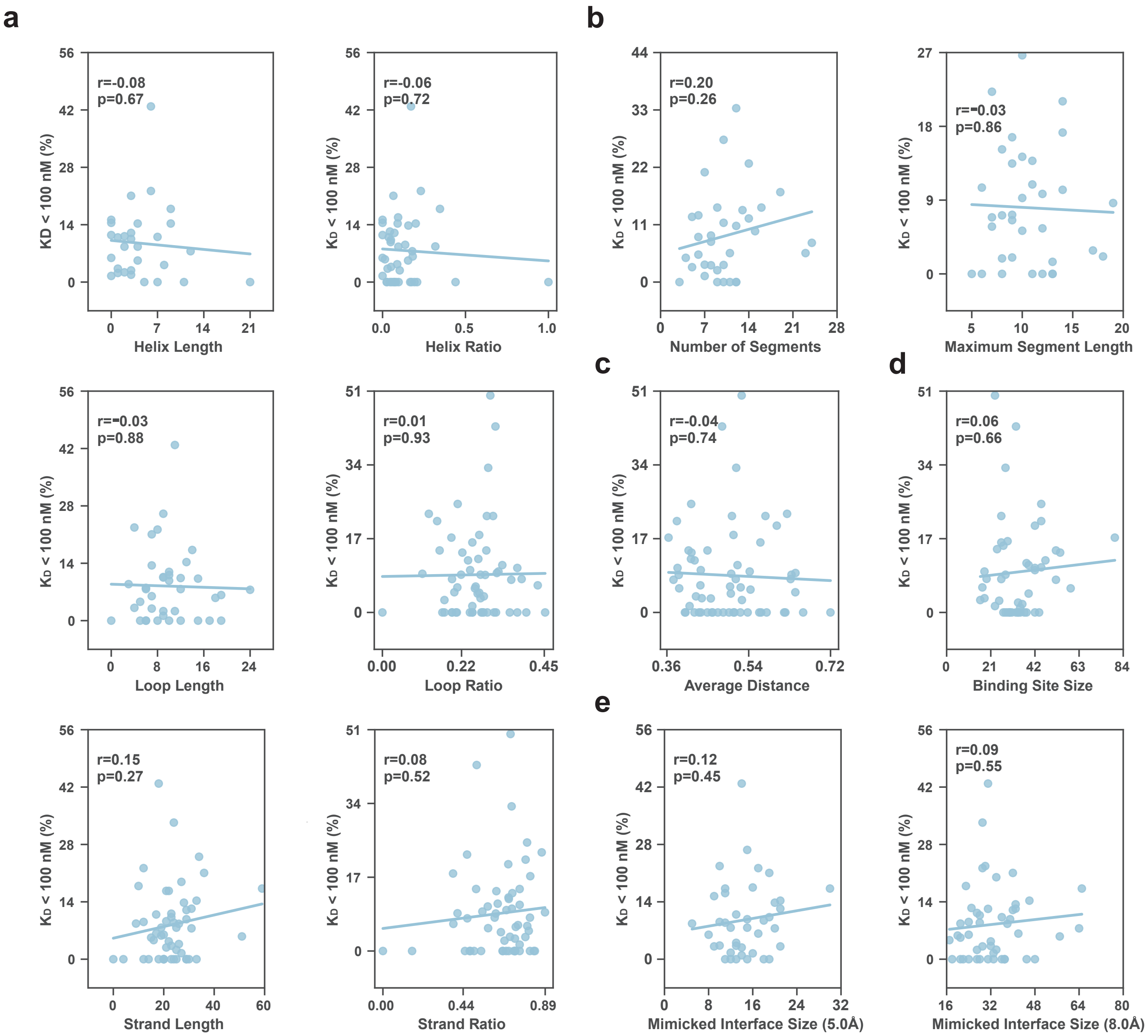

**Supplementary Figure 5. Correlation between success rates ($K_D < 100$ nM) and statistical properties on the reference interface for mimicry.** **(a)** Correlation between success rates and secondary structure compositions of reference binder. **Length** is defined as number of residues involved in the secondary structures, which is further normalized by the number of residues on the interface to obtain **Ratio**. **(b)** Correlation between success rates and number of segments on the reference interface of the binder, as well as the maximum length of segments. **(c)** Correlation between success rates and averge distances between residues on the reference interface of the binder. **(d)** Correlation between success rates and the size of binding site on the target protein. **(e)** Correlation between success rates and the number of residues on the reference interface, involving both the target protein and the binder.

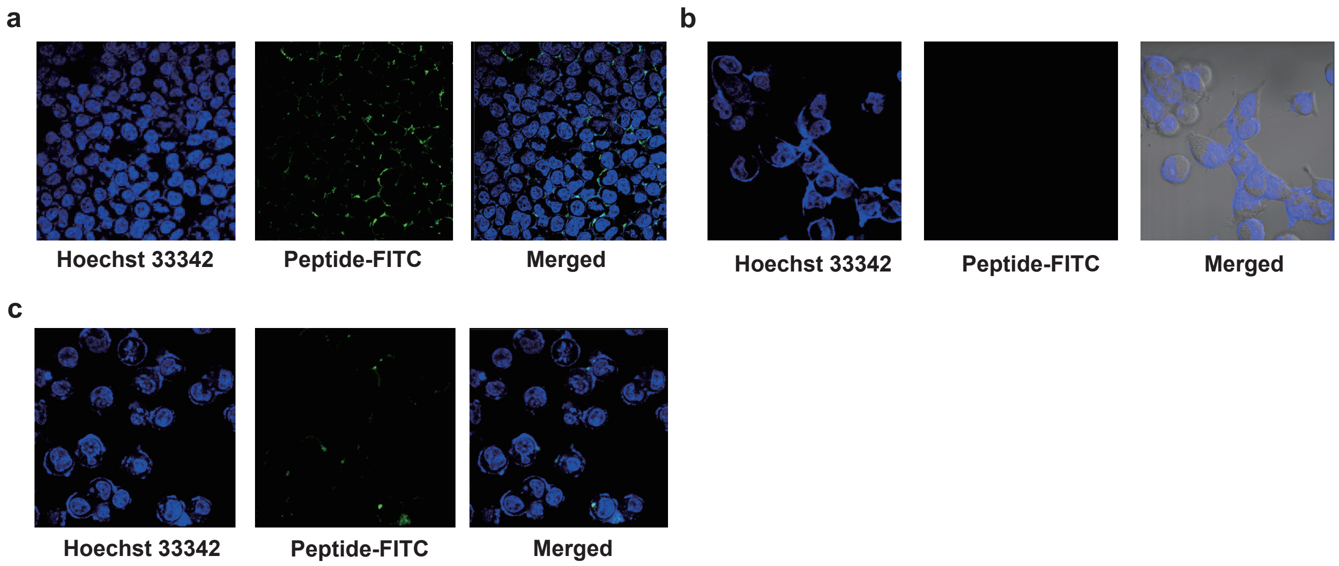

**Supplementary Figure 6.** **a**, Confocal image of the FITC labeled peptide RIWAYRKFMD binding towards PD-L1 negative cells 293T. **b**, Confocal image of the FITC labeled peptide NPDGAISWDVLQ binding towards CD38 negative cells 293T. **c**, Confocal image of the FITC labeled peptide DAIKDSEFENWL binding towards HER2 negative cells 293T.

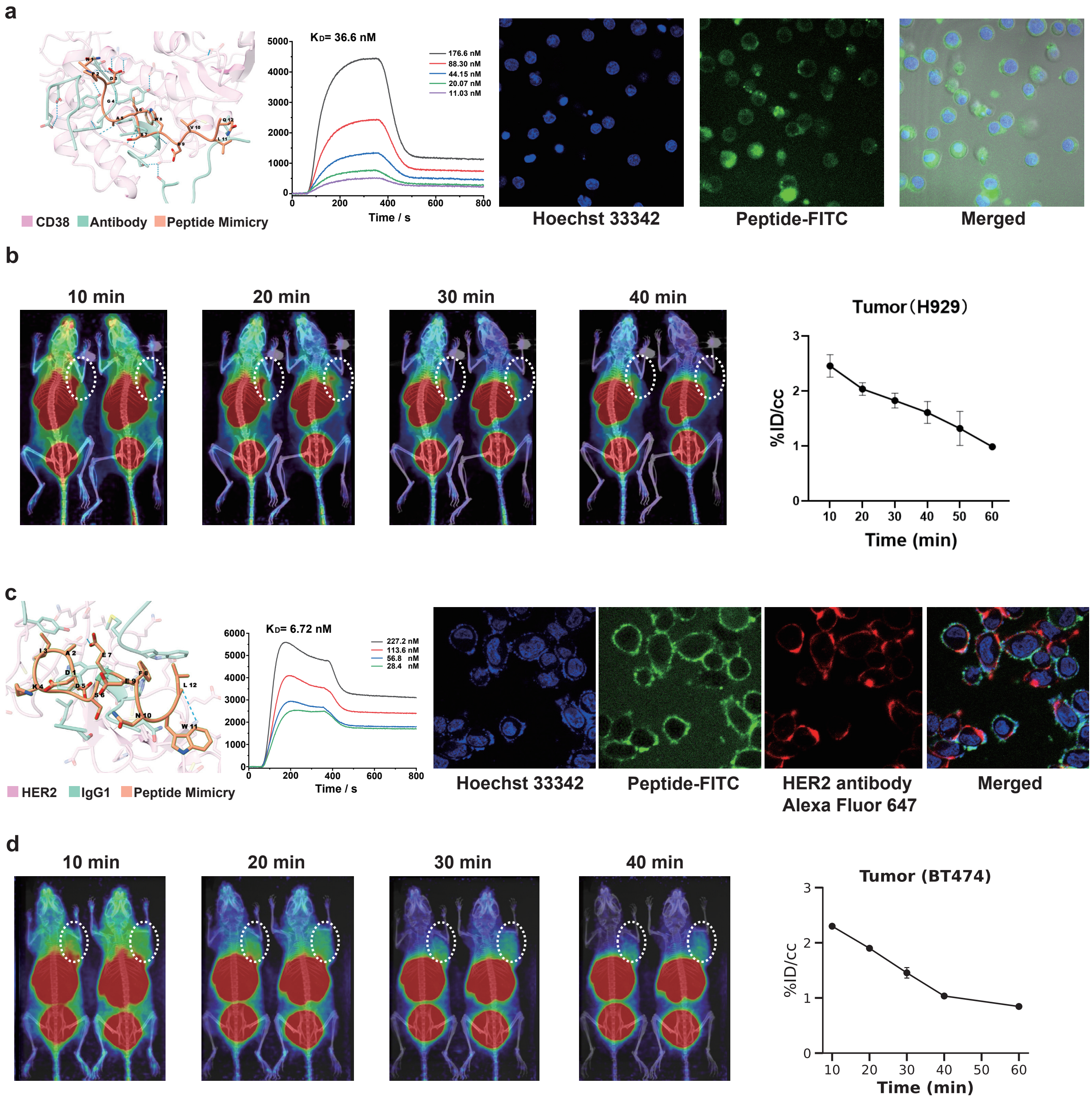

**Supplementary Figure 7.** **a**, The binding affinity of CD38 targeting peptide NPDGAISWDVLQ was analyzed by SPRi and confocal imaging on Ramos cells (CD38 positive) *in vitro*. Scale bar: 20 $\mu$m. **b**, Micro PET imaging of $^{68}$Ga–DOTA-NPDGAISWDVLQ in mice with CD38-positive H929 tumors xenografts. **c**, The binding affinity of HER2 targeting peptide DAIKDSEFENWL was analyzed by SPRi and confocal imaging on SKBR-3 cells (HER2 positive) *in vitro*. Scale bar: 20 $\mu$m. **d**, Micro PET imaging of $^{68}$Ga–DOTA-DAIKDSEFENWL in mice with HER2-positive SKBR-3 tumors xenografts.

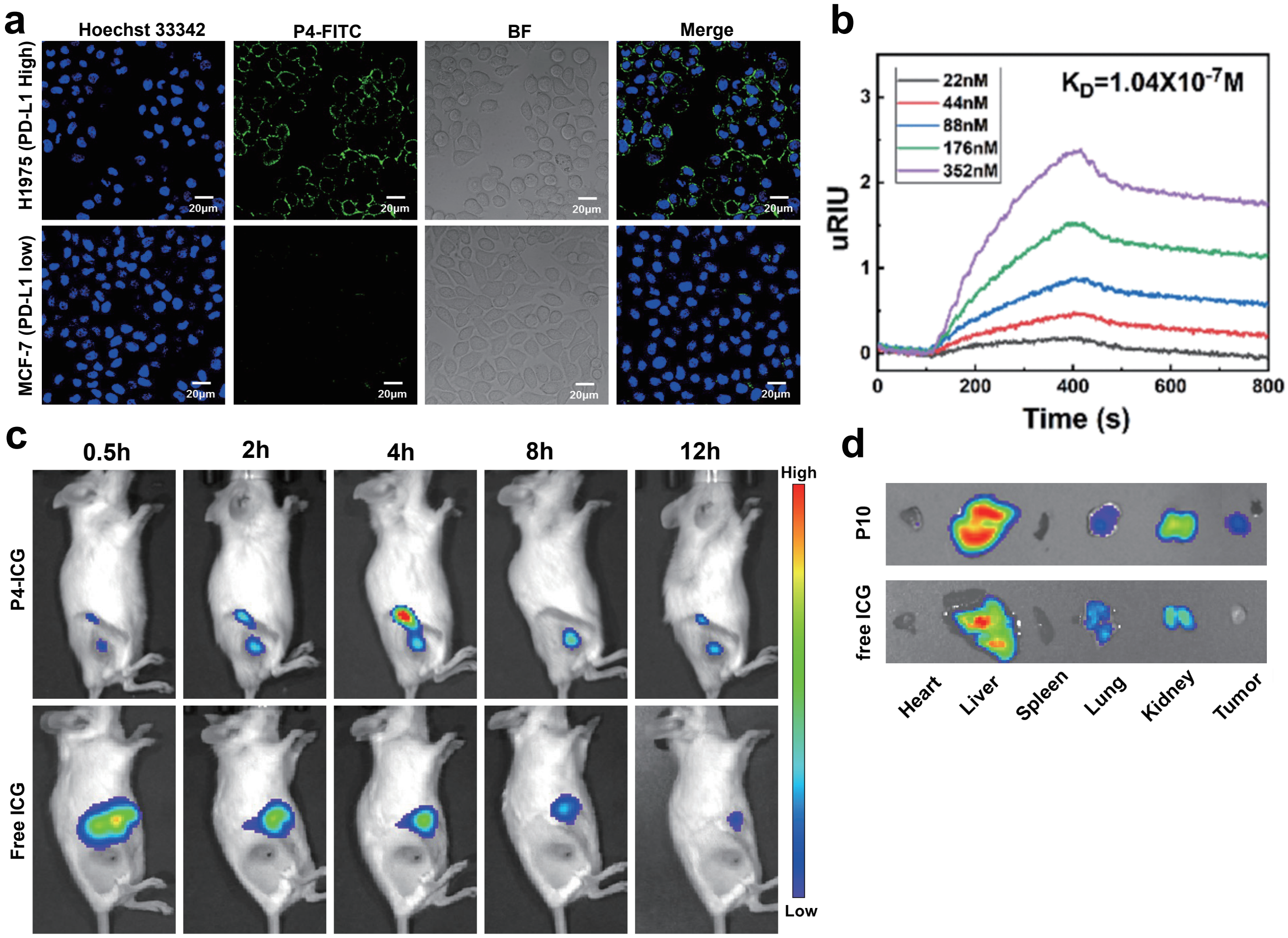

**Supplementary Figure 8. Affinity and specificity evaluation of PDL1 peptide probes.** **(a)**, Confocal imaging of the selected peptide binding to H1975 and MCF-7 cells. Scale bar: 20 $\mu$m. **(b)**, SPRi binding curves of the selected peptide with the PD-L1 protein. **(c)**, Fluorescence imaging of the selected peptide probe in mice with H1975 xenografts tumors. **(d)**, The representative fluorescence images of the main dissected organs at 12h after intravenous injection.

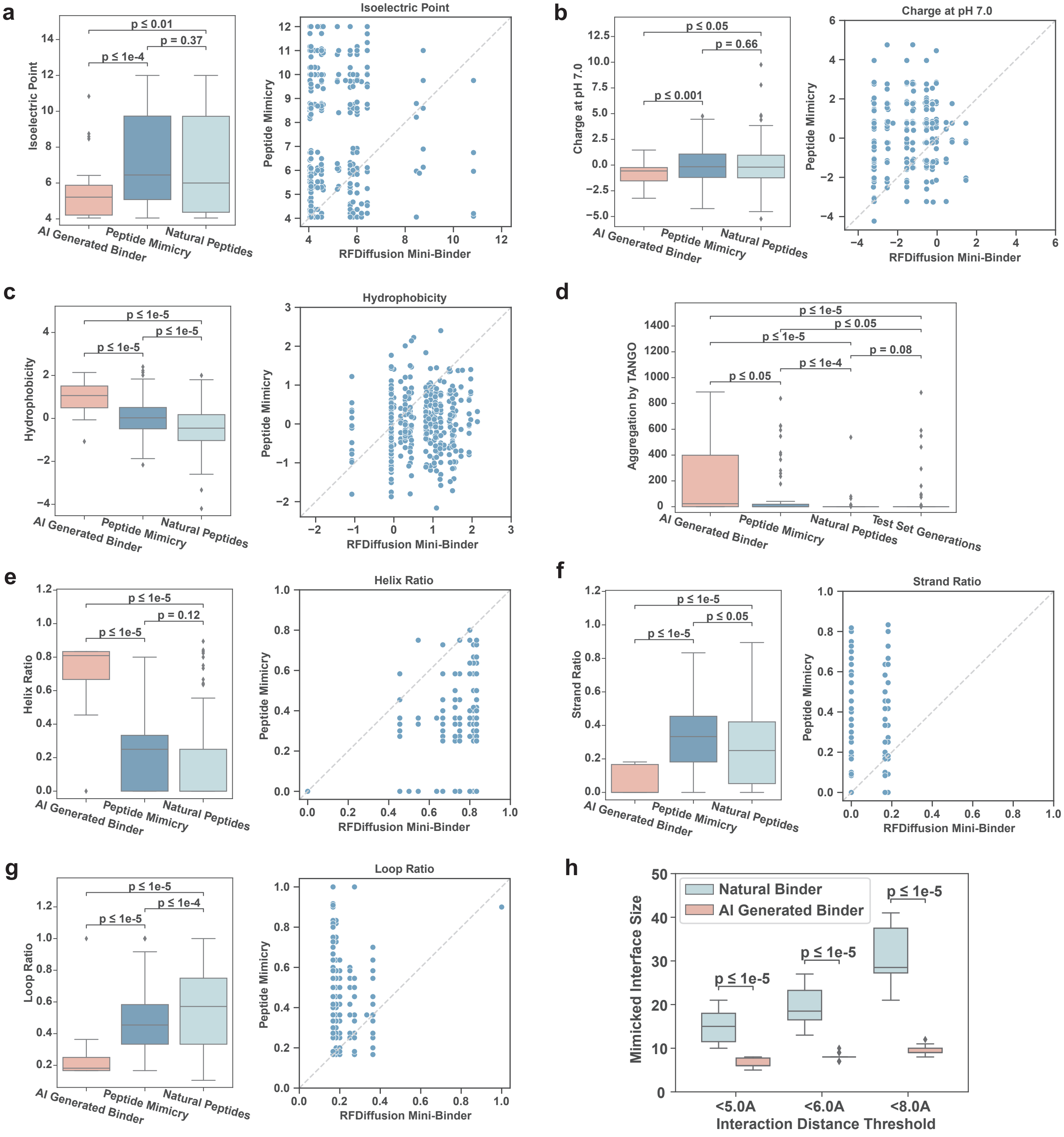

**Supplementary Figure 9. Comparison of physiochemical properties and secondary structures across peptides from various sources, including mini-binders generated by RFDiffusion, peptide mimicries from our algorithm, and natural peptides. (a-c)** Isoelectric point, charge at pH 7.0, and hydrophobicity distributions of peptides from different sources. The left parts in each panel show the overall distribution of the property values, while the right parts illustrate the changes in these properties when comparing the peptide mimicries to their reference mini-binders from RFDiffusion. **(d)** The comparison of aggregation propensity predicted by TANGO[6] across peptides from different sources. "Test set generations" refer to peptides generated on the test set by our model. **(e-g)** Comparison of secondary structure ratios in peptides from different sources. The left parts in each panel show the distribution of secondary structure ratios, and the right parts depict changes in these ratios when mimicking the mini-binders from RFDiffusion with our algorithm. **(h)** Comparison of interface sizes among different reference binders.

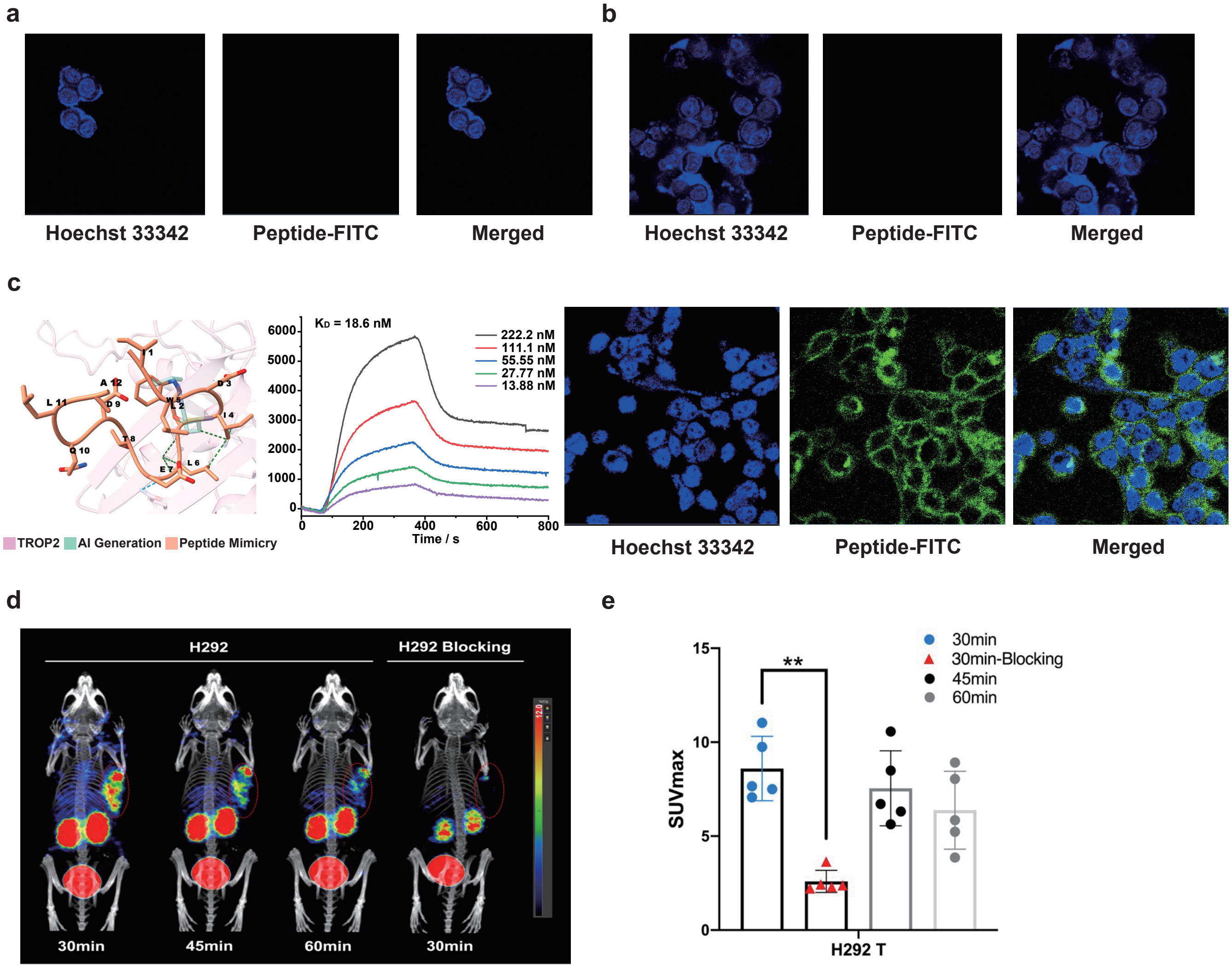

**Supplementary Figure 10.** **a-b**, The confocal imaging analyses for the TROP2 peptide binders PDMWTEAQLRFL and ILDIWLETDQLA on TROP2 negative cells 293T. **c**, Affinity and specificity evaluation of the selected TROP2-binding peptide ILDIWLETDQLA. SPRi binding curves of the peptide and the TROP2 protein. Confocal imaging of the binding of the peptide to BxPC3 cells. Scale bar: 20 $\mu$m. **d**, Micro PET images of $^{68}$Ga–DOTA-ILDIWLETDQLA in mice with H292 tumors xenografts and the H929-blocked group (n=3). **e**. Changes in tumor tissue SUVmax at 30 min, 45 min, and 60 min after tail vein injection of $^{68}$Ga–DOTA-ILDIWLETDQLA.

#### Supplementary Tables

**Supplementary Table 1.** PDB IDs of binding complexes used for mimicry design (PD-L1).

| Target | Complex PDB ID | Target Chains | Binder Chains | Binder Category |
|---|---|---|---|---|
| PD-L1 | 3bik | A | B | Natural Binder |
| PD-L1 | 3sbw | C | A | Natural Binder |
| PD-L1 | 4zqk | A | B | Natural Binder |
| PD-L1 | 5ggt | A | H,L | Antibody |
| PD-L1 | 5grj | A | H,L | Antibody |
| PD-L1 | 5ius | C | A | Natural Binder |
| PD-L1 | 5jds | A | B | Nanobody |
| PD-L1 | 5x8l | A | K,F | Antibody |
| PD-L1 | 5x8m | A | B,C | Antibody |
| PD-L1 | 5xj4 | A | H,L | Antibody |
| PD-L1 | 5xxy | A | H,L | Antibody |
| PD-L1 | 7c88 | M | H,L | Antibody |
| PD-L1 | 7czd | B | A | Nanobody |
| PD-L1 | 7tps | B | A | Natural Binder |
| PD-L1 | 7xad | A | C | Natural Binder |
| PD-L1 | 7yds | A | B,C | Antibody |
| PD-L1 | 8aok | A | B | Nanobody |
| PD-L1 | 8aom | A | V | Nanobody |

**Supplementary Table 2.** PDB IDs of binding complexes used for mimicry design (BCMA).

| Target | Complex PDB ID | Target Chains | Binder Chains | Binder Category |
|---|---|---|---|---|
| BCMA | 1oqd | K | A | Natural Binder |
| BCMA | 1xu2 | R | A | Natural Binder |
| BCMA | 4zfo | F | H,L | Antibody |
| BCMA | 6j7w | C | A | Nanobody |
| BCMA | 8hxq | C | A | Nanobody |
| BCMA | 8hxr | D | B | Nanobody |

**Supplementary Table 3.** PDB IDs of binding complexes used for mimicry design (CD38).

| Target | Complex PDB ID | Target Chains | Binder Chains | Binder Category |
|---|---|---|---|---|
| CD38 | 3raj | A | H,L | Antibody |
| CD38 | 4cmh | A | B,C | Antibody |
| CD38 | 5f1k | A | C | Nanobody |
| CD38 | 5f1o | A | B | Nanobody |
| CD38 | 5f21 | A | B | Nanobody |
| CD38 | 7dha | A | B,C | Antibody |
| CD38 | 7duo | B | H,L | Antibody |
| CD38 | 7vke | A | B | Nanobody |
| CD38 | 8byu | A | H,L | Antibody |
| CD38 | 8il3 | C | A,B | Antibody |

**Supplementary Table 4.** PDB IDs of binding complexes used for mimicry design (HER2).

| Target | Complex PDB ID | Target Chains | Binder Chains | Binder Category |
|---|---|---|---|---|
| HER2 | 1n8z | C | A,B | Antibody |
| HER2 | 1s78 | A | C,D | Antibody |
| HER2 | 3be1 | A | H,L | Antibody |
| HER2 | 3n85 | A | L,H | Antibody |
| HER2 | 3wlw | A | H,L | Antibody |
| HER2 | 3wsq | A | L,H | Antibody |
| HER2 | 4nnd | F | A | Natural Binder |
| HER2 | 5k33 | C | A | Antibody |
| HER2 | 5kwg | C | A | Antibody |
| HER2 | 5my6 | A | B | Nanobody |
| HER2 | 5o4g | C | A,B | Antibody |
| HER2 | 6att | A | H,L | Antibody |
| HER2 | 6bgt | C | A,B | Antibody |
| HER2 | 6j71 | A | B | Antibody |
| HER2 | 7mn6 | B | A,H | Natural Binder |
| HER2 | 8ffj | X | J,Q | Antibody |
| HER2 | 8ffj | X | I | Antibody |
| HER2 | 8hgo | B | A,C | Natural Binder |
| HER2 | 8pwh | E | A,B | Antibody |
| HER2 | 8pwh | E | C,D | Antibody |

**Supplementary Table 5.** PDB IDs of binding complexes used for mimicry design (CD4).

| Target | Complex PDB ID | Target Chains | Binder Chains | Binder Category |
|---|---|---|---|---|
| CD4 | 1jl4 | D | A,B,C | Natural Binder |
| CD4 | 3jcb | D | A | Natural Binder |
| CD4 | 3lqa | C | G,H,L | Natural Binder + Antibody |
| CD4 | 3o2d | A | L,H | Antibody |
| CD4 | 3s4s | G | A,B | Natural Binder |
| CD4 | 3t0e | E | A,B | Natural Binder |
| CD4 | 4q6i | C | A,B | Antibody |
| CD4 | 5cay | B | G | Natural Binder |
| CD4 | 6l1y | C | A | Natural Binder |
| CD4 | 6opn | C | A | Natural Binder |
| CD4 | 7t0r | A | G | Natural Binder |
| CD4 | 7t0r | A | L,H | Antibody |

**Supplementary Table 6.** Manual epitope selection on CD38 and TROP2 for peptide mimicry design without reference complexes. The epitope definitions follow the format: <Chain ID>:<Start Amino Acid><Start Residue Number>-<End Amino Acid><End Residue Number>. Non-continuous residues are separated with a comma. #Generation denotes number of candidates generated by RFDiffusion. #H-Sampling denotes times of resampling hotspots for RFDiffusion.

| Target | PDB ID | Eptiope Definitions | #Generation | #H-Sampling |
|---|---|---|---|---|
| CD38 | 3RAJ | A:P118-S295 | 12,000 | 600 |
| TROP2 | 7E5M | A:D171, A:R178, A:G241-P250 | 2,000 | 100 |
| TROP2 | 7E5M | A:Q237-Q252 | 2,000 | 100 |
| TROP2 | 7E5M | A:C34-K72 | 2,000 | 100 |
| TROP2 | 7E5M | A:L179-H187,A:Q252-Y260 | 2,000 | 100 |
| TROP2 | 7E5M | A:D146-R178 | 2,000 | 100 |
| TROP2 | 7E5M | A:V43-D65 | 2,000 | 100 |

**Supplementary Table 7.** Hyperparameters of PepMimic in sequence-structure codesign.

| Name | Value | Description |
|---|---|---|
| | | Variational AutoEncoder |
| embed size | 128 | Size of embeddings for residue types. |
| hidden size | 128 | Size of hidden layers. |
| $h$ | 8 | Size of the latent variable for residue types in the sequences. |
| layers | 3 | Number of layers. |
| mask ratio | 25% | Ratio of masked residues during encoding. |
| $\lambda_1$ | 0.3 | The weight of KL divergence on the sequence. |
| $\lambda_2$ | 0.5 | The weight of KL divergence on the structure. |
| $\lambda_{CA}$ | 1.0 | The weight of $C_\alpha$ loss in $\mathcal{L}_{aux}$. |
| $\lambda_{bond}$ | 1.0 | The weight of bond loss in $\mathcal{L}_{aux}$. |
| $\lambda_{angle}$ | 0.5 | The weight of side-chain dihedral angle loss in $\mathcal{L}_{aux}$. |
| | | Latent Diffusion Model |
| hidden size | 128 | Size of hidden layers in the denoising network. |
| layers | 3 | Number of layers. |
| steps | 100 | Number of diffusion steps. |
| RBF kernel | 32 | Number of Radial Basis Function kernels. |
| RBF cutoff | 3.0 | Distance cutoff for RBF kernels (in standard latent space). |
| | | Latent Interface Encoder |
| hidden size | 128 | Size of hidden layers in the encoder. |
| output size | 32 | Size of output, namely the interface embedding. |
| layers | 3 | Number of layers. |
| RBF kernel | 32 | Number of Radial Basis Function kernels. |
| RBF cutoff | 20.0 | Distance cutoff for RBF kernels (in unnormalized data space). |

**17.** I Loshchilov. Decoupled weight decay regularization. *arXiv preprint arXiv:1711.05101*, 2017.

**18.** Alexander Quinn Nichol and Prafulla Dhariwal. Improved denoising diffusion probabilistic models. In *International Conference on Machine Learning*, pages 8162–8171. PMLR, 2021.

**19.** Joost Schymkowitz, Jesper Borg, Francois Stricher, Robby Nys, Frederic Rousseau, and Luis Serrano. The foldx web server: an online force field. *Nucleic acids research*, 33(suppl_2):W382–W388, 2005.

**20.** Peter Vanhee, Almer M van der Sloot, Erik Verschueren, Luis Serrano, Frederic Rousseau, and Joost Schymkowitz. Computational design of peptide ligands. *Trends in biotechnology*, 29(5):231–239, 2011.

**21.** Joseph L Watson, David Juergens, Nathaniel R Bennett, Brian L Trippe, Jason Yim, Helen E Eisenach, Woody Ahern, Andrew J Borst, Robert J Ragotte, Lukas F Milles, et al. De novo design of protein structure and function with rfdiffusion. *Nature*, 620(7976):1089–1100, 2023.

**22.** Minkai Xu, Lantao Yu, Yang Song, Chence Shi, Stefano Ermon, and Jian Tang. Geodiff: A geometric diffusion model for molecular conformation generation. In *International Conference on Learning Representations*, 2022.
